# Dual-mode tethered intramolecular agonist-dependent mechanoactivation of the adhesion GPCR ADGRG1/GPR56

**DOI:** 10.1101/2025.09.16.676693

**Authors:** Chaoyu Fu, Cheng Quan, Lina Zielke, Yuan Weng, Ying Zhang, Gaojie Song, Tobias Langenhan, Jie Yan

## Abstract

Mechanical stimuli instruct cardinal cellular decisions pertaining to cell fate, proliferation, morphology, and movement^1,2^. How adhesion G protein-coupled receptors (aGPCRs), a large family of mechanosensors with more than 30 members in humans^3–5^, transduce mechanical cues into metabotropic commands has been a matter of debate due to the lack of suitable approaches to analyze receptor activation during mechanotransduction in live cells^6^. A central question is whether aGPCR activation strictly requires mechanical dissociation of the autoproteolyzed N- and C-terminal fragment (NTF and CTF) complex to expose the tethered intramolecular agonist (TIA; ‘*Stachel*’). Here, we investigated human ADGRG1/GPR56 (G1), an aGPCR with roles in brain development^7^, skeletal muscle^8^, and platelet function^9^, to study the events during aGPCR mechanotransduction by monitoring receptor dissociation and G protein recruitment in real time during cell migration and substrate deformation. Magnetic-tweezer measurements at a physiologically relevant loading rate showed that isolated G1 GAIN domains can partially unfold and refold at forces below those required for NTF-CTF dissociation. In cells, G1 dissociation occurred at retracting fibers during cell migration^10^ over an adhesive substrate through its native extracellular matrix ligand, tissue transglutaminase 2 (TG2), and was enhanced by cell stretching. Simultaneous live recording of G protein recruitment and pharmacological assays during mechanical stimulation of G1 reveal a dual-mode activation mechanism: mechanical force elicits robust metabotropic signaling without requiring NTF dissociation and generates an intracellular response commensurate with the fully dissociated receptor. Both signaling modes depend on an intact *Stachel* element^11,12^ of the receptor under our experimental conditions. Our findings determine mechanotransduction principles governing aGPCR activation at the cell-ECM interface. We reveal that G1 does not function as a simple binary switch, but rather as a highly sophisticated mechanosensor that dynamically encodes localized extracellular tension into distinct, highly scalable metabotropic outputs.

---

aGPCRs are complex cell surface molecules with extended extracellular regions (ECR) that contain receptor-specific adhesion motifs for interaction with cellular and matricellular ligands^5^. The ECRs are mounted on the archetypical heptahelical transmembrane domain (7TMD) of all GPCRs, while their intracellular regions (ICR) potentially provide anchoring to the cytoskeleton^4,5,13,14^. After biosynthesis many aGPCRs self-cleave into N-terminal (NTF) and C-terminal fragments (CTF) and form non-covalently associated complexes at the cell surface^15–18^. This remarkable post-translational feature is permitted through the presence of a GPCR autoproteolysis-inducing (GAIN) domain, the hallmark domain of all aGPCRs, which is located in the ECR near the membrane^18^. Receptor self-cleavage occurs at a GPCR proteolysis site (GPS) within the GAIN domain, which divides the GAIN domain itself into a larger N-terminal and a smaller C-terminal portion, the latter of which corresponds to the most C-terminal β-strand of the GAIN domain^18^. Apart from its structural role within the GAIN domain, this element stimulates receptor activity by interacting with the 7TMD as a tethered/intramolecular agonist (TIA; also termed *Stachel*). Ample pharmacological^11,12^, structural^19–22^ and *in vivo* evidence^9,23–25^ attests to this activation paradigm across the aGPCR family.

The molecular structure of aGPCRs lends itself to the activation by mechanical forces, e.g. during neuronal mechanosensation^6,26,27^, neural development^28^, myelination^29^, immune cell function^24,30,31^ or the control of surfactant levels in the lung^25^. The ECRs of individual aGPCRs sample the presence of local cognate adhesive ligands outside the receptor expressing cells, which laterally anchor them in the membrane^32^, and receive mechanical forces transmitted onto the receptors by relative movement between receptor and ligand. Incoming mechanical stimuli are thought to dissociate the NTF-CTF structure of autoproteolysed aGPCRs^28^ thereby exposing the *Stachel* to allow its interaction with the 7TMD (dissociation model) as exemplified for ADGRD1/Gpr133^33^, ADGRE2/EMR2^31^, ADGRE5/CD97^24,30^, ADGRF5/Gpr116^25^, G1^9^, and ADGRG6/Gpr126^29^.

Other studies demonstrated that, despite suppression of aGPCR self-cleavage via mutagenesis, receptor function can be retained *in vitro*^34^ and *in vivo*^27,35,36^ dependent on the integrity of the *Stachel* core sequence even though NTF-CTF separation is not feasible (non-dissociation model)^11,21,23,37,38^. This may involve intrinsic flexibility of the GAIN domain, allowing *Stachel*-7TMD engagement while the NTF and CTF remain associated^39,40^. In addition, single-molecule measurements indicated that separation of cleaved GAIN domains, and thus, NTF-CTF separation of a cleaved aGPCR, requires mechanical forces above a threshold defined by the non-covalent network of hydrophobic interactions within the autoproteolyzed domains^18^. E.g. G1, ADGRL1/LPHN1, and ADGRL3/LPHN3 dissociate at physiologically relevant forces of approximately 10-20 pN (refs. ^41,42^) after partial unfolding of the subdomain B of the GAIN domain, which occurs around 5-10 pN for both cleavable and non-cleavable receptor versions. Collectively, these findings imply that mechanical forces can stimulate aGPCR activity prior to receptor dissociation and transduce graded force stimuli before and after receptor dissociation, respectively. How aGPCR self-cleavage, dissociation, and activation are related to each other has been hampered by the lack of approaches to apply physiologically relevant mechanical forces to aGPCRs, independently observe the impact on their constituent NTF and CTF, and to simultaneously record their metabotropic output.

In order to assess the relationship between ligand-dependent mechanical stimulation of aGPCRs through physiologically relevant forces, GAIN domain autoproteolysis, receptor dissociation and intramolecular agonism, we chose the human aGPCR ADGRG1/Gpr56 (G1) as a model (Fig. 1a). G1 has a wide spectrum of physiological tasks in neural development^7^, immune^43^ and platelet^9^ functions, skeletal muscle homeostasis^8^, and relevance for the emergence and metastasis of neoplasias^44^. G1 is a minimalist aGPCR containing only pentraxin and laminin/neurexin/sex hormone-binding globulin-like (PLL) and GAIN domains in its ECR^45^ (Fig. 1a), which interacts with multiple ligands^44,46^. G1 is autoproteolysed^47^, its agonistic *Stachel* agonist has been defined and is required for receptor activation *in vitro*^12^ and *in vivo*^9^. Further, the cryo-EM structure of the active conformation of the G1-CTF has been obtained recently^20^. Finally, mechano-activated G1 couples to Gα_13_, and controls a RhoA-dependent pathway that governs cytoskeleton reorganization^9,12,44,47^.

**Fig. 1.**
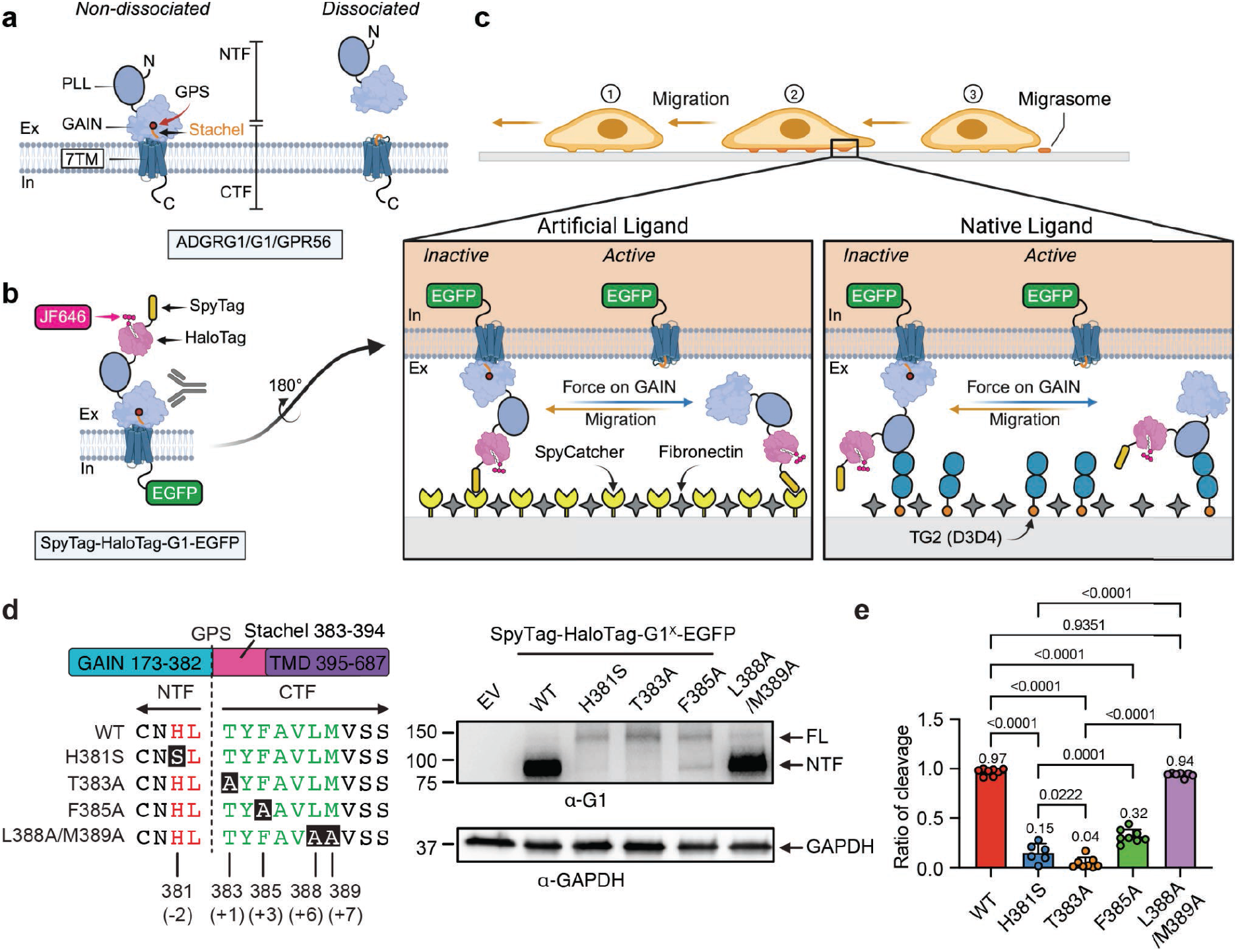
Methods for Real-Time Monitoring of GAIN Domain Dissociation and aGPCR Activation. **(a)** G1 receptor structure: PLL and GAIN domains in the ECD, a 7TMD, and an ICD. GPS cleavage produces NTF and CTF, which remain associated. Activation requires the intramolecular *Stachel* agonist and can occur via ligand or mechanical stimuli. **(b)** G1 receptor design for ‘migration force assay’: extracellular SpyTag and HaloTags enable adhesion (via SpyCatcher) and JF646 labeling, with an intracellular EGFP fusion. a-G1 targets the GAIN domain. **(c)** ‘Migration force assay’ setup: SpyTag-HaloTag-G1-EGFP adheres to SpyCatcher+FN-coated surfaces (left) or TG2 (D3D4)+FN-coated surfaces (right), visualizing receptor dissociation during cell movement. **(d)** G1 GPS sequence, autoproteolysis site, and signaling-impacting mutations (H381S, T383A, F385A, L388A/M389A) (left). Western blot analysis of COS-7 cells expressing SpyTag-HaloTag-G1 variants (G1^X^ corresponds to GPS mutations) (right). **(e)** Quantification of Western blot cleavage ratios for indicated G1 variants (summary for SpyTag-HaloTag-G1^x^-EGFP, SpyTag-G1^x^-EGFP in Ext. Data Fig. 3c and SpyTag-SNAPTag2-G1^x^-EGFP in Ext. Data Fig. 4i). Data (n ≥ 6 replicates for all groups) shown in a scatter plot with bar (data points plotted as mean ± SD); analyzed via an ordinary one-way ANOVA with Tukey’s multiple comparisons test. P values indicated.

## RESULTS

### G1 receptor design for migration-based mechanical activation by artificial or native adhesive ligands

To investigate ligand- and mechano-dependent aGPCR activation, we engineered a human G1 variant by inserting a HaloTag at the N-terminus of the PLL domain, which enables covalent fluorescent tracking of the G1-NTF in live cells via the addition of the organic dye JF646. Concomitantly, the receptor C-terminus was fused to EGFP to visualize the CTF. To facilitate faithful force transmission, an extracellular SpyTag was appended to the NTF, forming a stable covalent bond upon engagement with surface-coated SpyCatcher^48^ (Fig. 1b), serving as an artificial ligand interface for migrating COS-7 cells expressing the final SpyTag-HaloTag-G1-EGFP protein layout (Fig. 1c). For physiological comparisons, surfaces were alternatively coated with the D3D4 domains of tissue transglutaminase 2 (TG2) (Fig. 1c), a native endogenous ligand known to bind the G1-NTF (refs. ^44,49^).

We validated the autoproteolytic integrity of the SpyTag-HaloTag-G1-EGFP construct in COS-7 cells, a model lacking endogenous G1 expression (Fig. 1d). Furthermore, we generated a panel of receptor variants with mutations located in subdomain B of the GAIN domain flanking the GPS (H381S, T383A, F385A, and L388A/M389A) (ref. ^45^) (Fig. 1d) to differentially perturb self-cleavage and *Stachel*-dependent activation. The H381S mutation prevents GAIN domain-mediated autoproteolysis while retaining an intact agonistic *Stachel* sequence^18,45,50,51^. Mutations T383A and F385A also impair receptor self-cleavage but additionally perturb the *Stachel* sequence^41,51,52^. In particular, F385 is a highly conserved *Stachel* residue that is critical for G1 activation^12^. Finally, the L388A/M389A double mutant retains autoproteolysis but severely impairs TIA-dependent receptor activity^37^. Western blot analyses confirmed proper autoproteolysis of the wild-type (WT) fusion protein, generating the expected ∼95 kDa highly N-glycosylated NTF fragment (ref. ^53^) (Fig. 1d). The mutant variants, however, exhibited profoundly reduced cleavage efficiencies, evidenced by prominent ∼150 kDa full-length bands (Fig. 1d). Specifically, cleavage efficiencies plummeted to 15 % (H381S), 4 % (T383A), and 32 % (F385A) relative to WT (Fig. 1d-e, Extended Data Fig. 3c, Extended Data Fig. 4i). The L388A/M389A double mutant, designed to impair intramolecular agonism while preserving autoproteolysis, retained a 94 % cleavage efficiency (Fig. 1d-e, Extended Data Fig. 3c, Extended Data Fig. 4i).

### A real-time assay to monitor migration force and ligand-dependent aGPCR dissociation

To dissect the mechanotransduction sequence, we developed two distinct paradigms: a ‘migration force assay’ and a ‘deformation force assay’. The migration force assay exploits the SpyTag-SpyCatcher covalent bond as a robust, stable force-bearing point during cell migration. This interaction mediates force transmission to the G1-ECR at physiological loading rates in the pN/s range^54,55^, sufficient to separate the self-cleaved GAIN domain and dissociate the NTF-CTF complex^15,16,39^. We previously determined via single-molecule magnetic tweezers that the isolated G1-GAIN domain dissociates at 10-20 pN (at 1.0 pN/s)^41^, while single-molecule atomic force microscopy (smAFM) captured dissociation at 100-160 pN (at 1,000-40,000 pN/s), which could be extrapolated into 20 pN at equivalent loading rates^56^. Both physiological force thresholds remain securely below the ∼1.9 nN rupture force of the SpyTag-SpyCatcher bond (ref. ^48^).

COS-7 cells expressing the cleavable or non-cleavable G1 variants (Fig. 1d) were seeded onto fibronectin (FN) or a mixture of FN and SpyCatcher (FN+Spy) coated onto rigid polydimethylsiloxane (PDMS) substrates (2 MPa) (Fig. 1c). By leveraging dual-color labeling (NTF: JF646, magenta; CTF: EGFP, green), intact non-dissociated receptors appeared black via an inverted overlay (for better channel recognition the color palette was inverted; thus NTF+CTF is false-colored in black). Taking advantage of the rear detachment phenomenon^10^ inherent to migrating cells, we continuously imaged the trailing edges for 30 minutes (at 1 frame/min) starting 3 hours post-seeding (Fig. 2a, black circles; SI video. 1). This strategy allowed us to unequivocally distinguish between intact aGPCR complexes localized in migrating cells or migrasomes^13^, which are vesicles formed by plasma membrane material upon breaking off retracting fibres during migration^57^, versus dissociated G1-NTFs left firmly adhered to the ligand-coated surface tracks (Supplementary Fig. 1a).

**Fig. 2.**
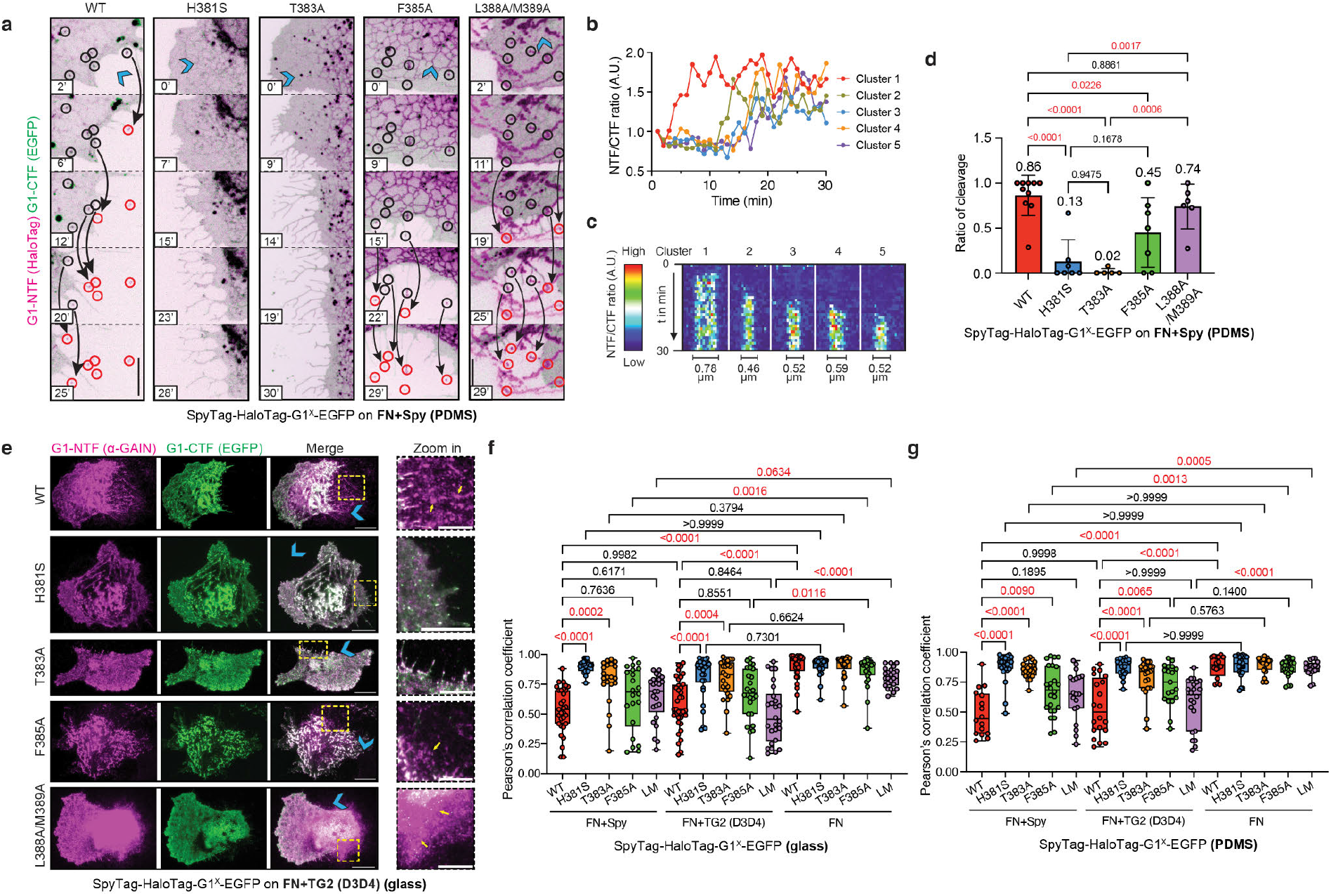
Visualizing of real-time GAIN domain dissociation during cell migration. **(a)** Visualization of SpyTag-HaloTag-G1^x^-EGFP dissociation in COS-7 cells on FN+Spy-coated PDMS surfaces. Frames (30’; 1 frame/min) show non-dissociated G1 (NTF [magenta]-CTF [green], black circles) and dissociated G1-NTF (magenta, red circles). Arrows indicate dissociation events; blue chevrons show migration direction. Scale bar: 10 µm. **(b)** Representative NTF/CTF fluorescence intensity traces and **(c)** Kymographs of individual G1 clusters over 30 min. The cluster size is shown below the spectrogram. **(d)** Cleavage ratio analysis (dissociated/total G1 clusters) for SpyTag-HaloTag-G1^x^-EGFP variants. Data (n≥5 replicates, 3 independent experiments) shown as mean ± SD scatter plots with bar; analyzed with an ordinary one-way ANOVA with Tukey’s multiple comparisons test. *P* values provided. **(e)** Immunostaining of G1-NTF (magenta) and fluorescence imaging of G1-CTF (green) in COS-7 cells migrating on FN+TG2 (D3D4)-coated glass surfaces. Blue chevrons indicate migration direction. Yellow arrows highlight dissociated NTFs (zoom-in). Scale bar: 20 µm; zoom-in: 10 µm. **(f-g)** Pearson’s correlation coefficient between G1-NTF and G1-CTF signals on glass **(f)** and PDMS **(g)** surfaces coated with FN+Spy, FN+TG2 (D3D4) and FN as measured in **(e)** and Ext. Data Fig. 1. Data (n > 20 (glass) and n > 15 (PDMS) replicates, 3 independent experiments) are shown in box-and-whisker plots (all data points plotted; median: horizontal line; boxes: 25th-75th percentiles; whiskers: min and max). Data were analyzed using an ordinary one-way ANOVA with Tukey’s multiple comparisons test.; P values provided.

In each image frame series, we assessed the dissociation ratio of individual receptor clusters based on their fluorescence signals. Prior to dissociation, G1 receptors were observed at the cell surface with a cluster, which had a mean projected area of 0.44 ± 0.17 µm^2^ (Supplementary Fig. 2a). During migration, JF646-labeled NTF-only fluorescent traces were progressively deposited in the migratory path, confirming active receptor dissociation (Fig. 2a, magenta circles; Supplementary Fig. 2b). The kinetics of GAIN domain separation were quantified by tracking the NTF/CTF fluorescence intensity ratio within fixed 0.8 × 0.8 µm regions of interest (ROIs) (Fig. 2b). Kymographic analyses of individual G1 clusters recorded an average NTF-CTF separation time of 6.2 ± 2.6 minutes (Fig. 2c, Supplementary Fig. 2c).

To rigorously confirm that dissociation is intrinsically dependent on GAIN domain autoproteolysis, we evaluated the cleavage-deficient receptor mutants. Consistent with our biochemical data (Fig. 1d-e), the NTF-CTF dissociation ratio for the cleavage-deficient variants were drastically reduced to 13 % for H381S, 2 % for T383A, and 45 % for F385A compared to WT (Fig. 2d). Conversely, the dissociation ratio of the cleavage-competent L388A/M389A variant was unaffected (Fig. 2d).

We further quantified GAIN domain dissociation via direct cell-surface fluorescence imaging following overnight migration (Fig. 2e, Extended Data Fig. 1a-e). To contrast the artificial SpyCatcher ligand against the native TG2 ligand, cells were mapped on FN+Spy, FN+TG2 (D3D4), or FN-only surfaces on both PDMS and glass substrates. Non-cleaved G1 on the thin membrane tubes was indicated by a white fluorescence signal. Dissociated G1-NTF ligated to the slide surface through SpyTag-SpyCatcher bonding or TG2 (D3D4)-G1-NTF interaction was indicated by a magenta fluorescence signal only (Fig. 2e, Extended Data Fig. 1a-e). Pearson’s correlation analysis confirmed that significant GAIN domain dissociation exclusively occurred in WT, F385A, and L388A/M389A variants migrating on either SpyCatcher or TG2-coated substrates (Fig. 2e-g). No dissociation was detected for the H381S or T383A mutations (Fig. 2e-g). Moreover, no dissociation was observed for any construct on FN-only surfaces lacking specific G1 ligands (Fig. 2f-g, Extended Data Fig. 1a-e). Both PDMS and glass conditions exhibited similar GAIN domain dissociation characteristics (Fig. 2e-g, Extended Data Fig. 1a-e).

To extend dissociation phenotype to another cellular background the migration force assay was independently replicated using Human Foreskin Fibroblasts (HFF). HFFs identically demonstrated that GAIN domain dissociation only occurred for the cleavage-competent WT and F385A variants strictly on FN+Spy surfaces, while the cleavage-deficient H381S and T383A mutants failed to dissociate (Supplementary Fig. 3a-b).

Collectively, these data establish the migration force assay as a highly quantitative tool for capturing aGPCR dissociation dynamics, confirming that physical fragment separation requires the obligate triad of ligand-receptor interaction, migration-induced force generation, and prior GAIN domain autoproteolysis.

### G1 dissociation occurs at retracting fibers within minutes after formation

Spatially, G1 dissociation is predominantly localized to the tips of retracting fibers oriented along the migratory axis at the cell’s trailing edge^10^ (Fig. 2). Time-lapse tracking in COS-7 cells revealed that these retracting fibers elongated at a characteristic rate of 0.77 ± 0.29 µm/min (Supplementary Fig. 2d-e). Continued cell migration eventually forced these fibers to snap, depositing migrasomes or retractosomes onto the substrate (Supplementary Fig. 1a)^58^.

Immunofluorescence analyses confirmed that both COS-7 and HFF models generated these structures, with retracting fibers containing both integrin α5 and actin filaments, whereas the resultant migrasomes retained integrin α5 but lacked actin (Supplementary Fig. 1b) (ref. ^59^). While G1 populated both the retracting fibers and migrasomes (Supplementary Fig. 1a-b), which is similar to ADGRE5/CD97 (ref. ^13^), active dissociation reliably occurred at a specific retracting fiber length of 4.56 ± 2.07 µm (Supplementary Fig. 2f), indicating G1 dissociation occurs within minutes of retraction fiber formation. Because maintaining a retracting actin-rich fiber requires forces similar to maintaining a filopodium, in the range of several pN at pulling rates in order of µm/min (ref. ^60^), this physiological tension perfectly matches the threshold required for the partial unfolding and subsequent dissociation of the G1-GAIN domain^41,42^. The structural integrity of retracting fibers and migrasomes remained unaltered in cells expressing the cleavage-deficient mutants, indicating that the failure to dissociate is a specific receptor property rather than a macroscopic cellular defect (Fig. 2e, Extended Data Fig. 1a-e, Supplementary Fig. 3a).

### Force-induced unfolding and dissociation of GAIN domains in G1 variants

Our migration force assay revealed distinct dissociation behaviors of the G1 variants. However, whether mutations near the GPS or within the *Stachel* sequence alter GAIN domain mechanical stability and thereby influence receptor signaling, remained unclear. We therefore examined isolated human G1 GAIN domains using a previously described magnetic-tweezers setup^41,61^ at a physiologically relevant loading rate of 2 pN/s (refs. ^54,55^). This approach enabled controlled force application and real-time measurement of changes in molecular extension (Fig. 3a). Among the mutants tested in the migration force assay, only H381S and T383A yielded sufficient purified protein for single-molecule measurements.

**Fig. 3.**
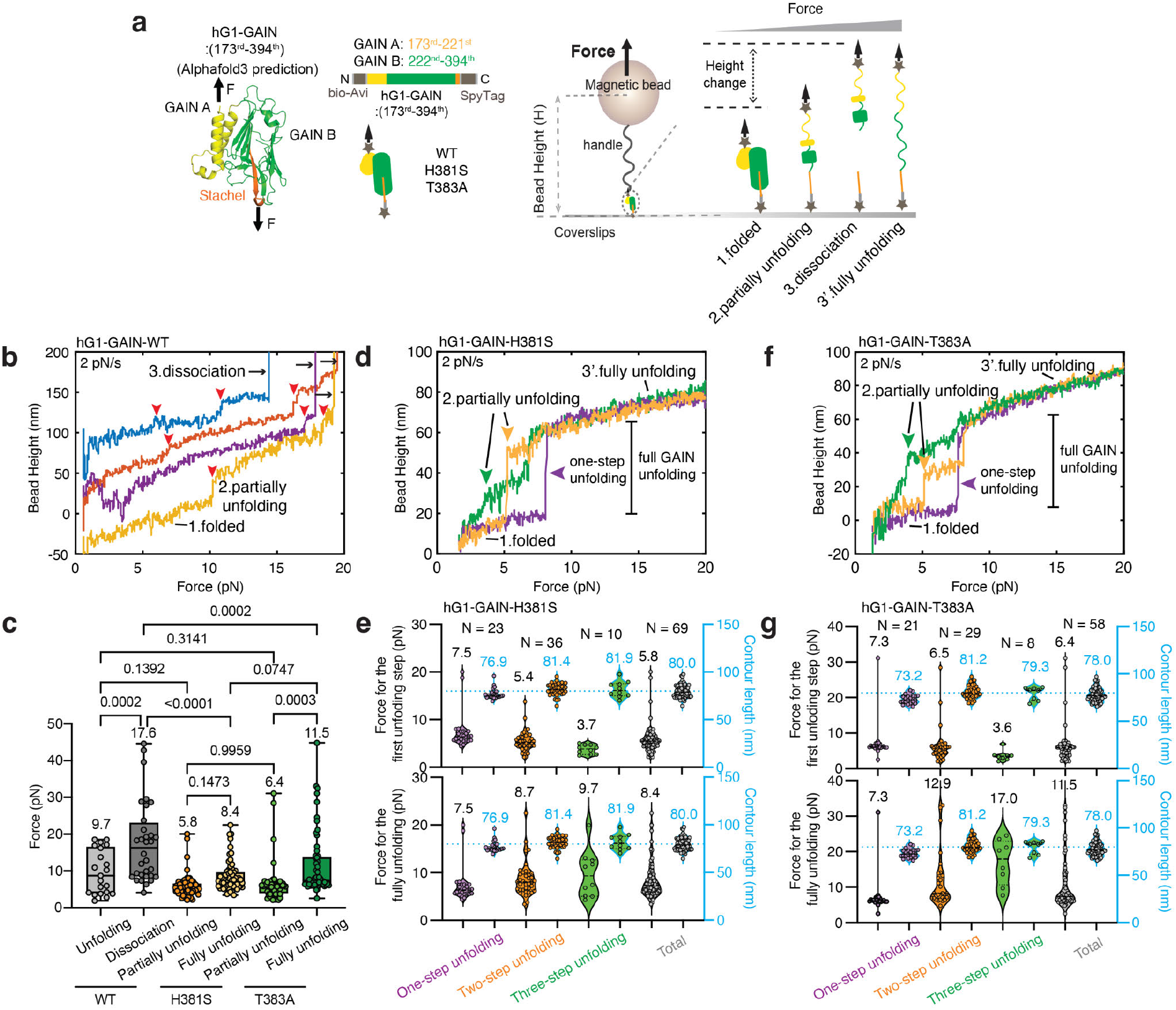
Force induced GAIN domain dissociation, partial or full unfolding in G1 variants. **(a)** Schematic of the single-molecule magnetic-tweezers assay used to characterize the mechanical stability of the human G1 GAIN domain (residues 173-394), together with its AlphaFold3-predicted structure. The AviTag- and SpyTag-flanked GAIN domain was tethered between a SpyCatcher-coated coverslip and a biotin-DNA-functionalized magnetic bead through Traptavidin. Force-induced structural changes were detected as changes in bead height. **(b)** Four representative force-height traces recorded for the WT G1 GAIN domain at a loading rate of 2 pN/s. Colored smoothed traces highlight unfolding transitions (red arrowheads) and NTF-CTF dissociation (black arrows). **(c)** Box plots of unfolding and dissociation forces for WT and mutant GAIN domains, showing the medians, interquartile ranges, and means. Each point represents a loading cycle. Independent tether numbers were 32 for WT, 8 for H381S and 5 for T383A; the corresponding cycle numbers were 32, 69 and 58. Cycles from the same tether represent repeated measurements. Experiments were performed in at least three independent replicates. Statistical significance was assessed by one-way ANOVA followed by Tukey’s multiple-comparisons test; *P* values are shown. **(d, f)** Three representative force-height traces for H381S (**d**) and T383A (**f**), illustrating single-, two-, and three-step unfolding pathways during repeated loading cycles. **(e, g)** Unfolding-force (black) and released-contour-length (blue) distributions obtained from all force-loading scans of H381S (**e**) and T383A (**g**), grouped according to the number of unfolding steps. Colored violin plots show the forces associated with the first and final unfolding transitions in each group. N denotes the number of loading cycles, mean values are indicated. Data were obtained from eight H381S tethers and five T383A tethers across at least three independent experiments.

Force-height traces from four independent G1 WT tethers showed one or more stepwise increases in molecular extension before NTF-CTF dissociation, consistent with varying degrees of partial unfolding (Fig. 3b). The first partial unfolding transition occurred at 9.7 ± 6.0 pN (mean ± SD; Fig. 3c). Consistent with our previous measurements^41^, G1 WT dissociated at 17.6 ± 10.9 pN at a loading rate of 2 pN/s (Fig. 3c).

By contrast, H381S and T383A showed no NTF-CTF dissociation and underwent repeated unfolding and refolding over multiple loading cycles (Fig. 3d, f). Between cycles, the force was reduced to 1 pN for 120 s to permit refolding. Three representative traces for each mutant revealed heterogeneous single-, two-, and three-step unfolding pathways, shown in purple, yellow, and green, respectively (Fig. 3d, f). We analyzed 69 and 58 force-loading scans for H381S and T383A, respectively, to determine the distributions of unfolding forces and step sizes for each event category (Extended Data Fig. 2a-d).

The number of residues released during each transition was estimated from the measured step sizes using a worm-like chain model with a persistence length of 0.8 nm. Summing the contour-length increments across all transitions yielded total released contour lengths of 80.0 ± 6.7 nm for H381S and 78.0 ± 7.4 nm for T383A (Fig. 3e, g), consistent with complete mechanical unfolding of the G1 GAIN domain (see **Methods**). Thus, although both mutations impair autoproteolysis, neither prevents the formation of a folded structure that resists pN forces during successive loading cycles.

The first unfolding transition occurred at 5.8 ± 3.0 pN for H381S (N = 69 scans) and 6.4 ± 5.1 pN for T383A (N = 58 scans) (Fig. 3e, g). Although the mean forces were lower than the first unfolding force measured for G1 WT (9.7 ± 6.0 pN), the differences were not statistically significant, suggesting these mutations do not substantially alter the mechanical stability of the GAIN structure. Based on the released contour lengths and known GAIN domain architecture, the first transition is consistent with partial unfolding of the predominantly β-stranded subdomain B, which contains the *Stachel* sequence (see **Methods**). The final unfolding transition occurred at 8.4 ± 4.0 pN for H381S and 11.5 ± 8.6 pN for T383A (Figure 3e, g), yielding contour lengths consistent with a fully unfolded conformation in which the *Stachel* sequence would be fully exposed. The mean forces associated with complete unfolding of both mutant GAIN domains were lower than that required for WT NTF-CTF dissociation at the same loading rate. These findings support the possibility that pN forces can partially or fully expose the *Stachel* sequence in the non-cleavable mutants.

Because WT partial unfolding occurred at lower forces than NTF-CTF dissociation, we also subjected WT to repeated low-force loading cycles. The force was increased to 15 pN at 2.0 ± 0.2 pN/s and then reduced to 1 pN for 120 s to permit refolding. Under these conditions, several WT GAIN-domain tethers underwent extensive partial unfolding at forces comparable to those observed for the mutants, without NTF-CTF dissociation (Extended Data Fig. 2e, purple curve). After several cycles, increasing the force beyond this range resulted in NTF-CTF dissociation (Extended Data Fig. 2e, blue curve). These findings demonstrate that WT GAIN can undergo repeated partial unfolding and refolding at forces below its dissociation threshold.

### G1 dissociation results in receptor activation and Gα_13_ recruitment in migrating cells

The non-dissociation model of aGPCR activation posits that receptor activation can occur in intact NTF-CTF receptor complexes, while dissociative signaling requires NTF release before CTF stimulation through tethered intramolecular agonism^3,4,62,63^. Both are relevant *in vivo*^5^. Because mechanical force can induce partial or full unfolding of the GAIN subdomain B independently of outright domain separation^40,41^, such changes potentially contribute to the activation of both autoproteolyzed and uncleaved aGPCRs^39^ in particular since several aGPCRs do not possess a self-cleaving GAIN domain by default^5^. Capitalizing on the migration force assay we directly tested the relationship between receptor dissociation and activation during mechanotransduction under physiological force application. Using a HaloTag-free construct (SpyTag-G1-EGFP) (Fig. 4a) expressed in COS-7 cells migrating on FN+Spy-coated glass (Fig. 4b), we utilized total internal reflection fluorescence microscopy (TIRFM) to monitor receptor mechanics. After 3 hours of seeding, immobilized G1 organized into distinct ridge-like arrays representing lateral membrane anchorage due to adhesion complex formation^32^ (Fig. 4c, d, SI Video. 2), termed ‘Spy adhesion sites’, indicating that G1 was covalently tethered to the SpyCatcher surface. Besides immobilized G1 ridges, highly mobile, non-tethered G1 puncta were also visible (SI Video. 2).

**Fig. 4.**
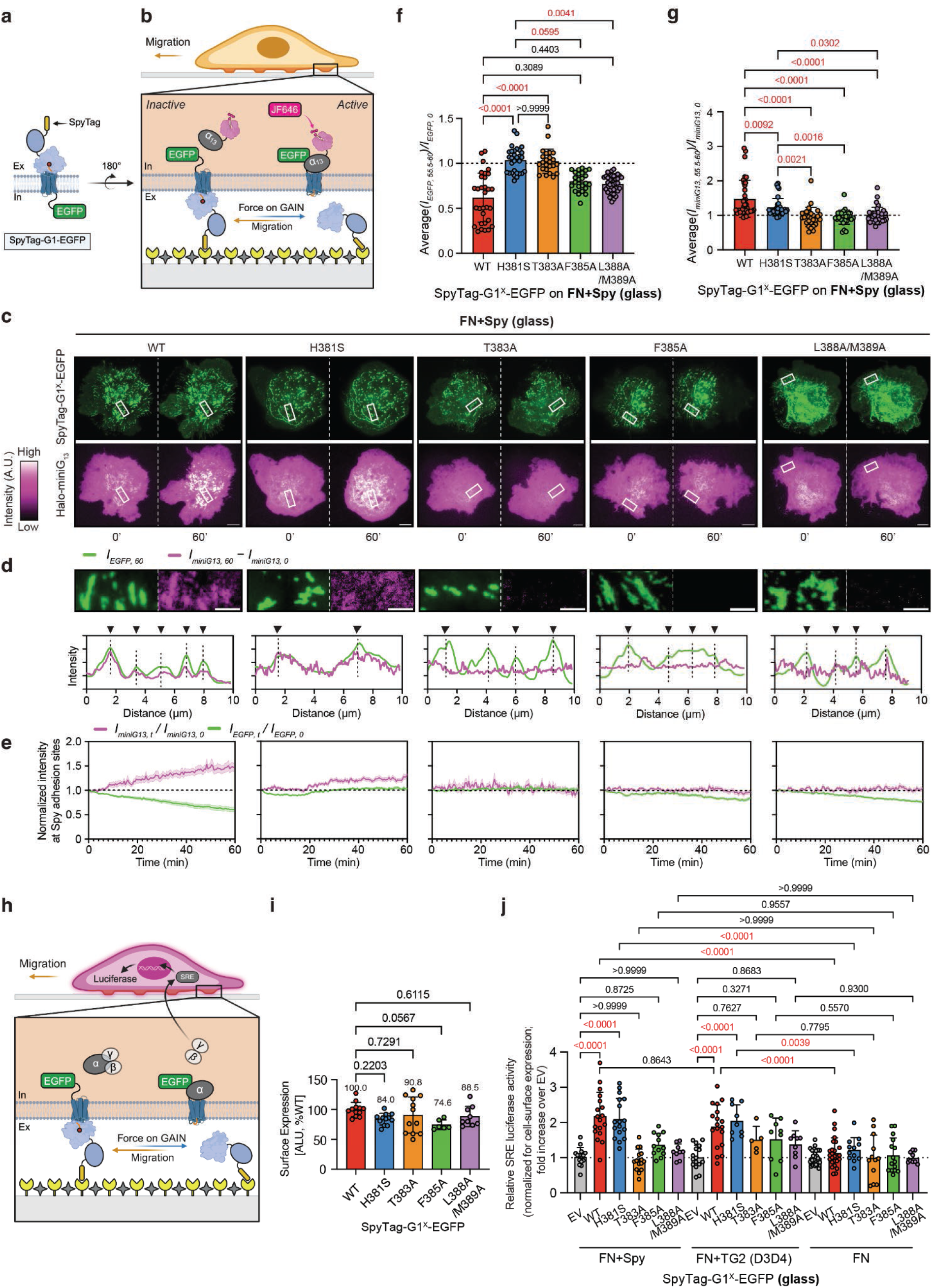
Dynamic mechanical activation of G1 during cell migration. **(a)** SpyTag-G1-EGFP receptor design for the ‘migration force assay’, combining G protein recruitment imaging. **(b)** Force-dependent GAIN domain dissociation at SpyCatcher sites activates G1, dynamically visualized via Halo-miniG13 recruitment and JF646 labeling in real-time using TIRFM. **(c)** TIRFM images of COS-7 cells expressing SpyTag-G1^x^-EGFP variants (green) migrating on FN+Spy-coated glass surfaces. HaloTag-miniG13 (magenta) is recruited to WT G1 and partially to the H381S mutant but not to *Stachel*-defective T383A, F385A and L388A/M389A variants. The net increase in signal at 60 minutes was shown in Ext. Data Fig. 3a. A zoomed-in white box (∼10 µm, shown in **(d)**) highlights the 60-minute time point. Scale bar: 10 µm. **(d)** Zoom-in region (top) and normalized intensity profiles (bottom) for G1 (green) and miniG13 (magenta) signal at Spy adhesion sites. MiniG13 intensity reflects net increases during the 60-minute interval. Dashed lines mark G1 signal peaks at 60 minutes. Scale bar: 3 µm. **(e)** Dynamic changes in normalized G1 intensity (I_EGFP,t_/I_EGFP,0_, green) and miniG13 intensity (I_miniG13,t_/I_miniG13,0_, magenta) at Spy adhesion sites over 60 minutes. Data are shown as line plots with mean ± SEM. **(f)** Average I_EGFP,60_/I_EGFP,0_ and **(g)** I_miniG13,60_/I_miniG13,0_ values from the final 5 minutes of imaging in **(e)**. Each point represents an adhesion site; at least 30 sites were sampled from at least six cells per group. Data are shown as scatter plots with mean ± SD. Statistical analysis: one-way ANOVA with Tukey’s test (P values shown). **(h)** G1 activation during the force migration assay was assessed via SRE-Luciferase activity. **(i)** Cell surface expression of SpyTag-G1^x^-EGFP variants. The receptor surface expression was quantified using ELISA (Methods). Data (n ≥ 6 replicates from 3 independent experiments) are shown as scatter plots with mean ± SD. **(j)** Relative luciferase activity for SpyTag-G1^x^-EGFP variants on FN+Spy, FN+TG2 (D3D4) or FN-coated glass surfaces during migration. Mutant constructs normalized for cell-surface expression from **(i)**. Dataset before normalization is shown in Ext. Data Fig. 3d. Data (n ≥ 5 replicates from 3 independent experiments) are shown as scatter plots with mean ± SD. Statistical analysis: one-way ANOVA with Tukey’s test (P values shown).

As these immobilized receptors sustained migration-dependent force transmission, we surmised that G1 could be mechanically activated there. Because G1 activation leads to engagement of Gα_13_ (refs. ^9,12,20^), we dynamically recorded real-time signal transduction by tracking the recruitment of a fluorescent HaloTag-miniG13 probe^64^ by JF646-labeling, a proximal readout of receptor conformations capable of engaging the engineered probe (Fig. 4b) under live conditions during cell migration with TIRF imaging (SI Video. 2). At Spy adhesion sites subjected to mechanotransduction, robust HaloTag-miniG13 recruitment was visualized (Fig. 4c, Extended Data Fig. 3a, SI Video. 2). MiniG13 fluorescence at Spy adhesion sites increased over time and, during prolonged imaging, subsequently declined (Supplementary Fig. 4a-b, SI Video. 3). These trajectories do not distinguish changes in receptor activity from probe redistribution, receptor trafficking or photobleaching^65^.

### The cleavage-deficient H381S variant retains signaling activity

Strikingly, dynamic TIRFM revealed that HaloTag-miniG13 was robustly recruited not only to the WT receptor but also to the cleavage-deficient H381S mutant (Fig. 4c, d, Extended Data Fig. 3a), which retains the NTF-CTF complex structure (Fig. 1d). In contrast, no miniG13 recruitment occurred for the *Stachel*-perturbed T383A, F385A, or L388A/M389A variants (Fig. 4c, Extended Data Fig. 3a). These results were confirmed in TIRF-ROI scans of SpyTag-G1-EGFP and JF646-HaloTag-miniG13 intensities (Fig. 4d).

To quantify this, we analyzed dynamic changes in normalized fluorescence intensities (I_EGFP_/I_EGFP,0_ (green) and I_miniG13_/I_miniG13,0_ (magenta)) across multiple Spy adhesion sites (>30 sites per condition) at 2 frame/min (Fig. 4e) (see Methods). For WT G1, we recorded a significant increase in I_miniG13_/I_miniG13,0_ coupled with a decrease in I_EGFP_/I_EGFP,0_, indicating rapid G protein recruitment outpacing the subsequent receptor internalization or out-diffusion of the shed CTF (Fig. 4e-g, SI Video. 2). Conversely, the cleavage-deficient H381S mutant exhibited a steady, moderate increase in I_miniG13_/I_miniG13,0_ while maintaining a constant I_EGFP_/I_EGFP,0_ level, demonstrating continuous activation with markedly reduced receptor dissociation (Fig. 4e-g, SI Video. 4). The T383A variant remained immobilized (constant I_EGFP_/I_EGFP,0_) but failed to activate (constant I_miniG13_/I_miniG13,0_) (Fig. 4e-g, SI Video. 5). The F385A and L388A/M389A mutants displayed no miniG13 recruitment despite undergoing partial or robust dissociation, respectively (decreasing I_EGFP_/I_EGFP,0_), confirming activation strictly requires an unperturbed *Stachel* (Fig. 4e-g, SI Video. 6-8).

Control experiments confirmed that this activation is purely force- and ligand-dependent: no miniG13 recruitment for any G1 variants occurred on FN-only surfaces (Extended Data Fig. 3b) or when utilizing SpyTag-deficient receptors on FN+Spy substrates (Supplementary Fig. 5a-c), supporting a requirement for receptor anchoring. Furthermore, arresting cell migration via treatment with 100 nM Dasatinib (Supplementary Fig. 6a), a Src kinase inhibitor^66^, thoroughly abolished both GAIN domain dissociation and miniG13 recruitment (Supplementary Fig. 6b-g), strongly supporting that active cellular traction forces are an absolute prerequisite for G1 signaling.

### Non-dissociative signalling elicits G1 activation comparable to dissociative signalling

To capture the cumulative metabotropic output of this mechanical activation, we utilized a serum response element (SRE)-luciferase reporter assay (Fig. 4h). Because standard cell-lysis and Western blot extraction efficiencies can be heavily biased by the differing solubilities of cleaved versus uncleaved GPCR fragments, we directly quantified the relative cell-surface expression of each intact G1 construct using an intact-cell ELISA (Fig. 4i). The measured SRE activity (Extended Data Fig. 3d) was subsequently normalized to these surface expression levels (Fig. 4j), ensuring that our quantitative functional comparisons were based strictly on equivalent plasma membrane receptor pools.

Following surface normalization, cells expressing WT G1 on FN+Spy surfaces displayed robustly elevated SRE activity compared to non-ligated FN-only conditions (Fig. 4j), which did not differ significantly from empty vector (EV) controls under the conditions of this assay(Fig. 4j). These results describe ligand- and force-dependent signaling in the present assay and do not exclude basal activity reported using other experimental systems. Remarkably, the SRE activation level of the cleavage-deficient H381S mutant on FN+Spy surfaces matched that of the fully dissociative WT receptor (Fig. 4j). This confirms again that, while the H381S mutant severely impacts autoproteolytic activity of the G1-GAIN domain (Fig. 1d-e), migration-dependent force exerted can non-dissoctively activate the receptor. As expected, the *Stachel*-deficient mutants (T383A, F385A, L388A/M389A) completely failed to activate during migration (Fig. 4j).

The migration force assay enabled us to recapitulate both receptor dissociation and activation, supporting an association between force-induced receptor dissociation and *Stachel*-dependent signaling. To verify the role of intramolecular agonism in this mechanotransduction, we applied the partial G1 antagonist dihydromunduletone (DHM)^67^. Treatment with 10 µM DHM entirely eradicated both the elevated SRE activity and the localized miniG13 recruitment for WT cells migrating on FN+Spy (Supplementary Fig. 7a-e). These findings demonstrate that forces generated during cell migration enable the *Stachel*-dependent mechanical activation of G1, which can be mitigated by DHM.

Previous studies reported that G1 activation reduces cell migration speed of neural progenitors^68^ and neurons^46^. We further measured the migration speed of COS-7 cells expressing the indicated G1 versions. Tracking cellular velocities revealed that COS-7 cells expressing signaling-competent receptors (WT and H381S) exhibited significantly reduced migration speeds on FN+Spy compared to FN surfaces (Extended Data Fig. 3e), whereas activation-deficient variants (T383A, F385A) displayed no such deceleration (Extended Data Fig. 3e), indicating that not mere adhesion forces between G1 and its ligands account for the speed reduction. Together, these normalized readouts prove that mechanical stimulation induces identical magnitudes of metabotropic signaling whether the GAIN domain physically sheds or undergoes non-dissociative conformational rearrangement.

In summary, these results demonstrate mechanical force activation of G1 through force-dependent GAIN domain dissociation and non-dissociative conformational GAIN domain changes. Interestingly, non-dissociative mechanical stimulation of G1 results in similar metabotropic signaling levels to stimuli that caused GAIN domain dissociation, and both modes require an intact *Stachel* sequence in our assays.

### Dual-mode mechanical activation of G1 via its native ligand TG2 during cell migration

To ensure the findings that utilize the SpyCatcher/SpyTag system represent physiological responses that can reconcile the interaction of G1 with one of its native adhesive ligands, we evaluated force transmission between G1 and TG2 (refs. ^44,49^), whose interaction with G1 regulates extracellular matrix remodeling and melanoma progression (ref. ^44^). Cells migrating on FN surfaces co-coated with the TG2 (D3D4) domain yielded WT and H381S activation profiles virtually identical to the FN+Spy condition (Fig. 4j), whereas the T383A, F385A, and L388A/M389A mutants remained functionally silent (Fig. 4j).

We then engineered a microcontact-printing array to directly visualize dynamic activation on native ligands, restricting biotinylated TG2 (D3D4) strictly to 12 μm neutravidin-coated circular regions separated by 10 μm FN-coated gaps (see Methods) (Fig. 5a). To visualize G1 dissociation and activation simultaneously, we utilized a SpyTag-SNAPTag2-G1-EGFP construct, in which the former HaloTag was replaced with a SNAPTag2^69^ (Fig. 5b), and fluorescently labeled it with tetramethylrhodamine (TMR). COS-7 cells coexpressing Spy-SNAP2-G1-EGFP and Halo-miniG13 were plated on TG2 (D3D4)-coated micropatterns (Fig. 5b). In TIRFM imaging, overlapping GFP and TMR signals (appearing yellow) were observed as circular adhesions of ∼12 µm in diameter, indicating G1 localization to TG2 (D3D4)-coated micropatterns (Fig. 5c-d, SI Video. 9-10). During migration across these micropatterns, TMR-labeled NTF traces were deposited by WT cells, confirming robust, native ligand-mediated dissociation (Fig. 5c, SI Video. 9-10). Crucially, Halo-miniG13 recruitment occurred exclusively within the geometric boundaries of the TG2 patterns, proving simultaneous dissociation and activation (Fig. 5e-g, SI Video. 9-10).

**Fig. 5.**
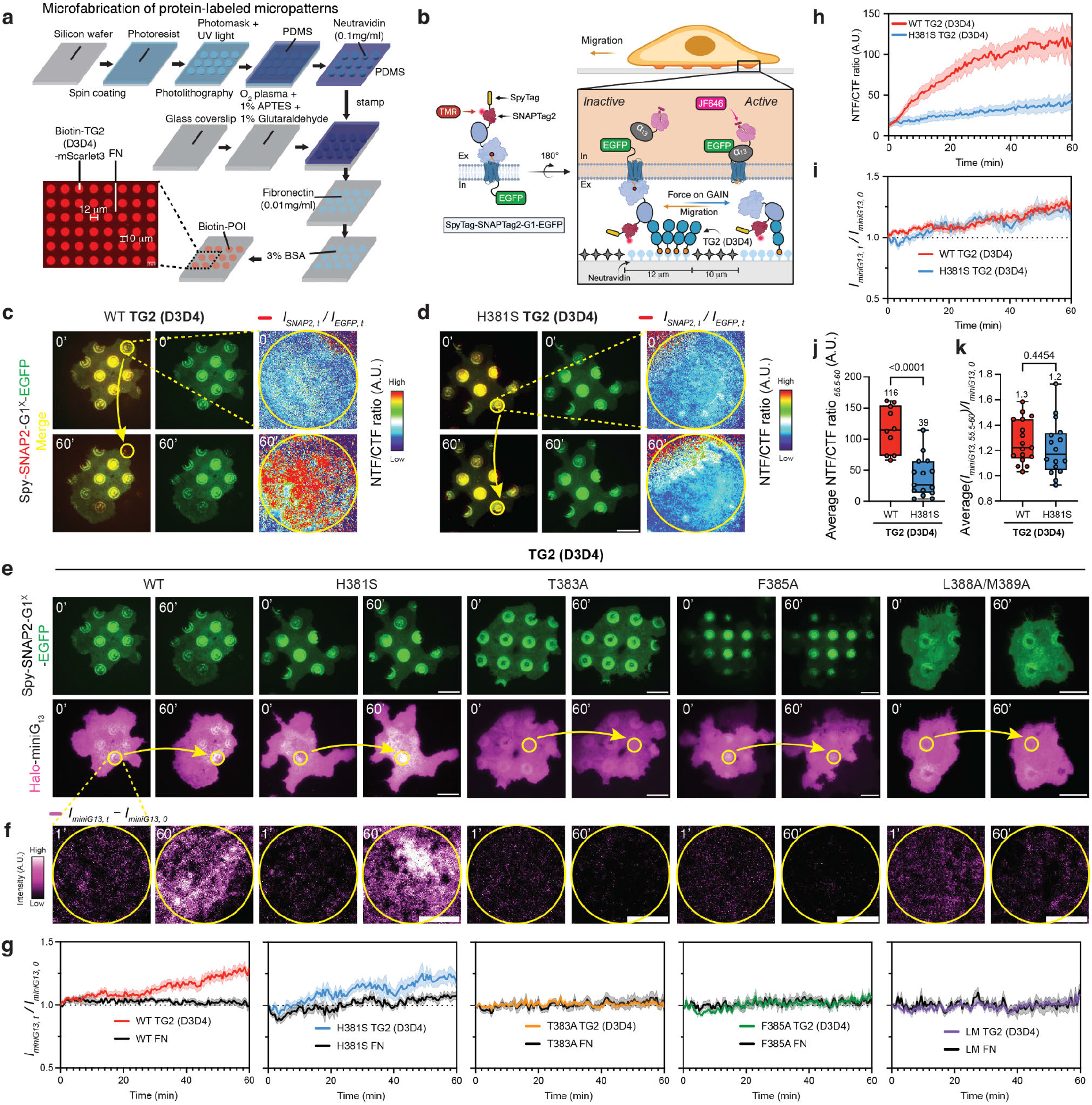
Dual-mode mechanical activation of G1 via its native ligand TG2 during cell migration. **(a)** Schematic illustration of the microfabrication of protein-labeled micropatterns. Circular microcontact-printed patterns of neutravidin (12 µm in diameter, 10 µm gap) were fabricated to bind biotin-labeled proteins of interest (POIs). Biotin-TG2 (D3D4)-mScarlet3 was used for visualization. FN was coated in the surrounding area to support cell migration. **(b)** Schematic of the SpyTag-SNAPTag2-G1-EGFP receptor design used in the ‘migration force assay’, integrating G1 dissociation and G protein recruitment imaging. Force-dependent GAIN domain dissociation at TG2 (D3D4)-coated micropatterns activates G1, which is visualized in real time by Halo-miniG13 recruitment and JF646 labeling using TIRFM. G1 dissociation is detected through the mislocalization of the NTF (TMR-labeled SNAPTag2) and CTF (EGFP) under TIRFM. **(c-d)** TIRFM images of COS-7 cells expressing SpyTag-SNAPTag2-G1^x^-EGFP variants (yellow: Merge of SNAPTag2, red, and EGFP, green) migrating on TG2 (D3D4)-coated micropatterns. Scale bar: 20 µm. The yellow circle (12 µm diameter) indicates the zoomed-in region showing G1 dissociation. The NTF/CTF fluorescence ratio reflects the extent of G1 dissociation. Scale bar: 5 µm. G1 dissociation occurs in WT but not in H381S. **(e)** TIRFM images of COS-7 cells expressing SpyTag-SNAPTag2-G1^x^-EGFP variants (green) migrating on TG2 (D3D4)-coated micropatterns. Scale bar: 20 µm. HaloTag-miniG13 (magenta) is recruited to both WT G1 and the H381S mutant, but not to *Stachel*-defective T383A, F385A and L388A/M389A variants. The yellow circle (12 µm diameter) indicates the zoomed-in region shown in **(f)**. **(f)** Zoom-in region showing miniG13 recruitment at TG2 (D3D4)-coated micropatterns. The miniG13 intensity represents the net increase during the 60-minute interval. Scale bar: 5 µm. **(g)** Time traces of dynamic changes in normalized miniG13 intensity (I_miniG13,t_/I_miniG13,0_, magenta) at TG2(D3D4)-coated micropatterns versus surrounding FN over 60 minutes for indicated SpyTag-SNAPTag2-G1^x^-EGFP variants. Data (n ≥ 10 replicates) are shown as line plots with mean ± SEM. **(h)** Time traces of NTF/CTF fluorescence intensity showing G1 adhesion dynamics at TG2(D3D4)-coated micropatterns over 60 minutes for WT and H381S. **(i)** Dynamic changes in normalized miniG13 intensity (I_miniG13,t_/I_miniG13,0_, magenta) at TG2(D3D4)-coated micropatterns over 60 minutes for WT and H381S. This dataset is also shown in **(g)**. **(j)** Average NTF/CTF ratio and **(k)** I_miniG13,t_/I_miniG13,0_ values from the final 5 minutes of time traces in **(h-i)**. Data (n ≥ 10 replicates) are shown as scatter plots with mean ± SD. Statistical analysis: Welch’s t-test (P values shown).

For the H381S mutant, we observed potent, spatially restricted miniG13 recruitment on the TG2 micropatterns, whereas the pronounced TMR/EGFP mislocalization observed for WT was not evident (Fig. 5d-g, SI Video. 10-11). This combination of retained signaling and reduced fragment separation is consistent with a contribution from non-dissociative activation.The *Stachel*-impaired variants (T383A, F385A, L388A/M389A) failed to activate on the TG2 patterns (Fig. 5e-g, SI Video. 12-14), phenocopying the SpyCatcher results.

To precisely quantify these coupled dynamics, we measured both the NTF/CTF fluorescence ratio, which was calculated as the intensity of the TMR-labeled NTF over the EGFP-labeled CTF, and the corresponding Halo-miniG13 intensity within each discrete TG2 (D3D4)-coated micropattern. During cellular transit, localized receptor dissociation, particularly at retracting fibers near the rear edge, drove a marked increase in the NTF/CTF ratio within the patterned areas for G1-WT cells. Conversely, no significant ratio change was detected for the cleavage-deficient G1-H381S variant (Fig. 5h, SI Video. 10-11). This divergence was further corroborated by direct cell-surface imaging following overnight migration; orthogonal staining with an anti-GAIN domain antibody confirmed robust receptor shedding in WT cells, but a substantially reduced dissociation in the H381S mutants (Supplementary Fig. 8a-b). Yet, the H381S mutant exhibited substantial receptor activation, generating elevated miniG13 intensities directly over the TG2 (D3D4) micropatterns that mirrored WT levels, in stark contrast to the constant value for the surrounding FN (Fig. 5i, SI Video. 10-11).

The ligand specificity and force-transmission capacity of this assay were validated through parallel controls. Negative control surfaces utilizing uncoated patterns yielded absolutely no G1 enrichment or activation, standing in stark contrast to the targeted engagement seen on the TG2 (D3D4) arrays (Supplementary Fig. 9a-b). Furthermore, as a positive control for load-bearing capacity, we replicated these experiments using the artificial covalent SpyCatcher linkage. Similar dissociation-associated changes and signaling responses were observed on SpyCatcher-coated patterns (Extended Data Fig. 4a-h, Supplementary Fig. 9a-b). These findings indicate that both anchoring systems can support receptor dissociation and activation under the tested conditions.

Crucially, our orthogonal multi-tag design (SNAP-Tag/EGFP for structural dissociation and Halo-miniG13 for metabotropic activation) permitted the simultaneous comparison of these two parameters within the same cell and patterned region. Quantitative cellular analysis revealed a stark decoupling of physical shedding from functional output: while the WT receptor dissociated at a roughly three-fold higher than the H381S mutant on TG2 (D3D4)-coated micropatterns, their corresponding levels of G protein activation were nearly identical (Fig. 5j-k). This identical uncoupling of dissociation and signaling was reproduced when force was transmitted via SpyCatcher-coated patterns (Extended Data Fig. 4g-h). These results definitively demonstrate that the markedly elevated physical dissociation of the WT receptor does not yield a proportionally higher G protein activation compared to the cleavage-deficient H381S variant. Consequently, the robust signaling profile of H381S cannot be attributed merely to an undetected dissociated fraction, strongly validating the existence of a highly efficient, non-dissociative mechanical activation pathway.

Overall, these findings demonstrate that spatially restricted mechanical tension, transmitted exclusively through the native ligand TG2, provides sufficient mechanical stability to drive both G1 dissociation and activation, and G1 signaling under mechanical force that does not require G1 dissociation, dispensing with the necessity for highly artificial tethers.

### G1 encodes a progressive increase in mechanical stimulation into increased signalling activity

To extend our findings to secondary scenarios where cells physiologically experience force, we modeled force transmission during macroscopic substrate deformation. In this ‘deformation force assay’, COS-7 cells expressing SpyTag-HaloTag-G1-EGFP were seeded onto flexible PDMS elastomers co-coated with FN+Spy and subjected to a 16 % uniaxial stretch for 1 hour (Fig. 6a, Supplementary Fig. 10a). This specific expansion range accurately models the 5-20 % extracellular matrix (ECM) distension characteristic of physiological processes and is widely utilized in mechanotransduction studies^70^. We hypothesized that cell migration and external stretch might act as additive mechanical stimuli, driving heightened GAIN domain dissociation and elevated G1 activation. Indeed, the combination of both active migration and external stretch significantly amplified GAIN domain dissociation (Fig. 6b-c). As expected, no dissociation occurred on FN-only surfaces lacking the requisite SpyCatcher anchor (Fig. 6b-c).

**Fig. 6.**
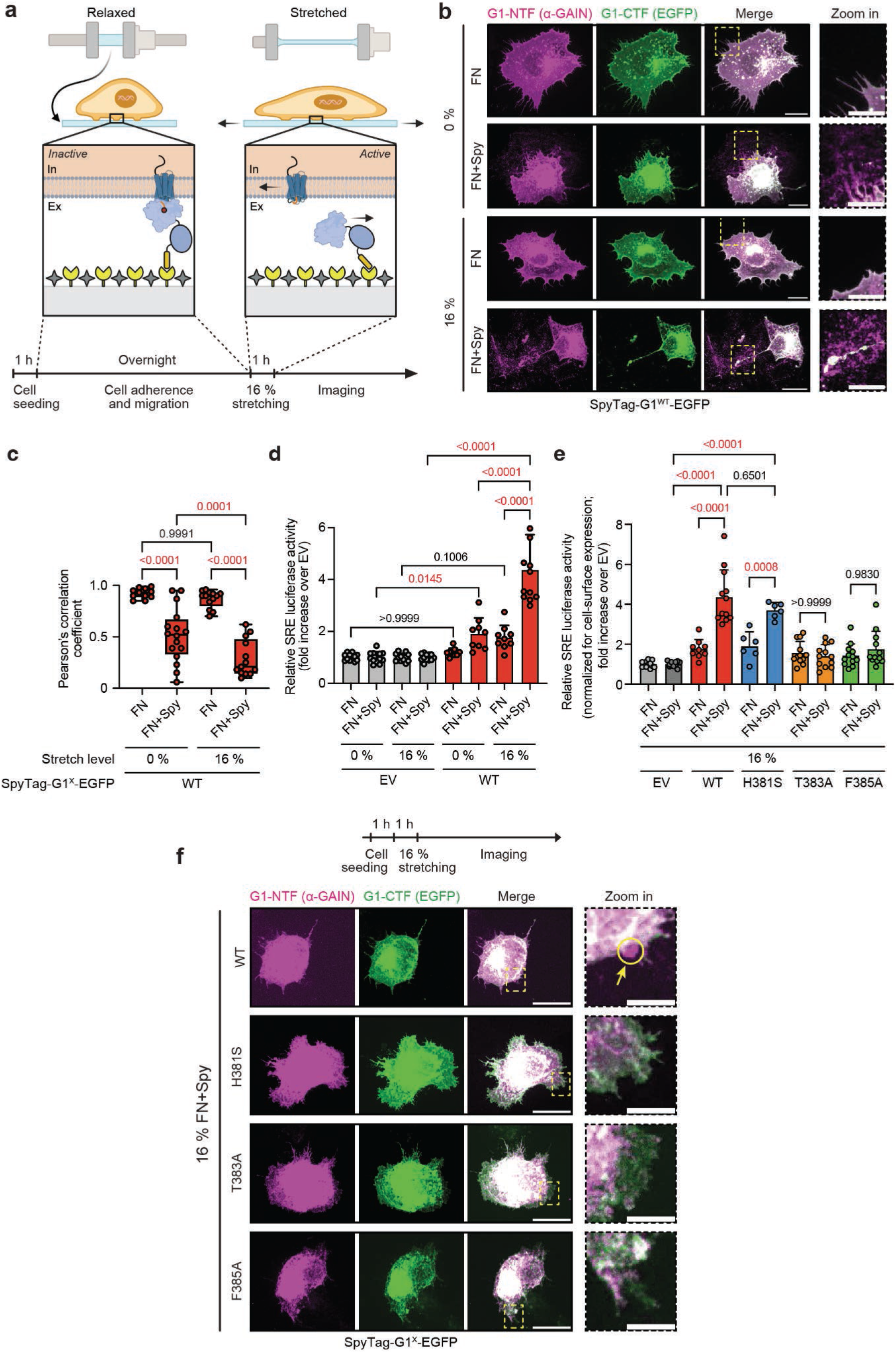
Effects of substrate stretching on G1 dissociation and signaling. **(a)** G1 dissociation and activation triggered by ECM stretching in the ‘deformation force assay’. **(b)** Immunostaining of COS-7 cells expressing SpyTag-G1^WT^-EGFP on FN or FN+Spy surfaces, under unstretched or 16 % stretched conditions. NTF (α-GAIN, magenta) and CTF (EGFP, green) are visualized. Scale bars: 20 µm (main) and 10 µm (zoom-in). **(c)** Correlation between NTF (α-GAIN) and CTF (EGFP) signals as measured in **(b)**. Pearson’s correlation coefficients were calculated. Data (n > 10 biological replicates from at least 3 independent experiments) are shown as box-and-whisker plots (all data points plotted; median: horizontal line; boxes: 25th-75th percentiles; whiskers: min and max). One-way ANOVA with Tukey’s test was performed, with P-values indicated. This dataset is also shown in Ext. Data Fig. 5b. **(d)** G1 activation in the ‘deformation force assay’ assessed by SRE-luciferase reporter assay under conditions in **(b)**. Empty vector (EV) served as a normalization control. Data (n > 6 replicates from 3 independent experiments) are shown as scatter plots with bars (mean ± SD). One-way ANOVA with Tukey’s test were performed; P-values are shown. This dataset is also shown in Ext. Data Fig. 5c. **(e)** Activation of G1 variants upon 16 % stretch. Data are presented as in **(d)**. Mutant constructs normalized for cell-surface expression from Fig 4i. This dataset is also shown in Ext. Data Fig. 5d. **(f)** Immunostaining of COS-7 cells expressing SpyTag-G1-EGFP variants on FN+Spy surfaces under 16% stretch. Cells were fixed post-seeding and stretching. Color scheme as in **(b)**. Yellow arrows in the zoom-in region indicate dissociated NTF. Scale bars: 20 µm (main) and 5 µm (zoom-in).

Consistently, WT G1-expressing cells mapped on FN+Spy substrates exhibited a highly significant amplification in SRE-luciferase receptor activation when subjected to the deformation force, compared to unstretched controls (Fig. 6d). Furthermore, baseline SRE-luciferase activity was intrinsically higher on FN+Spy surfaces compared to FN-only surfaces, reinforcing the absolute necessity of a stable mechanical linkage at the G1-NTF to enable force transduction (Fig. 1c, Fig. 6d, Supplementary Fig. 10b).

Remarkably, the H381S variant also registered robust, stretch-induced activation enhancements on FN+Spy-coated substrates compared to FN-only controls (Fig. 6e, Extended Data Fig. 5a-d). This behavior directly reinforces our core conclusion: a cleavage-deficient G1 receptor harboring an intact *Stachel* sequence is fully capable of non-dissociative mechanical activation.

The stretch-dependent escalation in G1 activation strongly implies that macroscopic ECM deformation accelerates structural GAIN domain dissociation. To isolate this variable and directly visualize stretch-induced shedding, cells were seeded for a brief 1-hour window to minimize baseline migration-dependent dissociation while permitting adequate cell spreading^71^, prior to applying the 16 % stretch for 1 hour. Post-stretch imaging confirmed that rapid GAIN domain dissociation occurred explicitly at the cell periphery of stretched WT G1 cells (Fig. 6f), proving that deformation-induced structural separation executes within a 1-hour timeframe.

Our data imply a graded mechanosensory mechanism: physiological tension initially induces activating conformational deformations within the intact receptor, while escalating mechanical stress breaches the structural threshold to forcibly dissociate the GAIN domain, unmasking the full signaling capacity of the GPCR.

## DISCUSSION

Resolving aGPCR mechanotransduction is essential for understanding receptor physiology^4,63^ and for pharmacological manipulation^72^. A major limitation has been the lack of methods that simultaneously resolve receptor dissociation and G protein coupling during mechanical stimulation. Physiologically defined stimuli have been studied only in a few models, including neuronal proprioceptive, acoustic and tactile sensory systems^23,26,27,73^, blood-borne^9,30^ or flow-exposed cells^24,74^, and the lung^25^. Direct measurement of force-dependent metabotropic signaling in vivo remains difficult, whereas in vitro assays often rely on mechanical stimulation regimes that compare poorly with physiological conditions^29^, or on artificial activation by antibodies^75–78^ or proteases^33,37,79^.

Here, we addressed this gap by applying mechanical stimuli to the human G1 receptor using cell migration and substrate deformation as force generators. Signaling of WT G1required self-cleavage and NTF-CTF dissociation, whereas the cleavage-deficient mutants^12,18,45,51,52^ allowed to distinguish dissociation from signaling. During force application, miniG13 recruitment and SRE activation showed that G1-H381S, which retains an intact TIA, remained signaling-competent, whereas TIA/*Stachel* mutants G1-T383A and G1-F385A, which showed reduced cleavage, and the cleavage-competent TIA/*Stachel* mutants G1-L388A/M389A were inactive, consistent with prior work in G1^9,12,20^ and other aGPCRs^19,23,37,38^. Thus, the native *Stachel* sequence is required for G1 mechanotransduction under our assay conditions.

In single-molecule magnetic-tweezer experiments, isolated H381S and T383A GAIN domains underwent repeated unfolding transitions without tether dissociation, with released contour lengths consistent with extensive unfolding (Fig. 3). WT GAIN domains also underwent repeated partial unfolding and refolding at forces below its dissociation threshold (Extended Data Fig. 2e). These data support a two-regime mechanotransduction model for G1 (Fig. 7). The high-force, binary signaling mode resembles the irreversible ‘one-and-done’ activation of Notch^80^ and the dissociation model of aGPCR activation^4,12^. At lower forces typical of retracting fibers during cell migration (ref.^60^), approximately 5-10 pN can partially unfold GAIN subdomain B (ref.^41^), expose the *Stachel*, and drive scaled, potentially reversible GAIN-7TMD coupling without fragment separation. Above the approximately 10-20 pN dissociation threshold (ref.^41^), NTF and CTF separate, fully uncapping the *Stachel* and producing full activation.

**Fig. 7.**
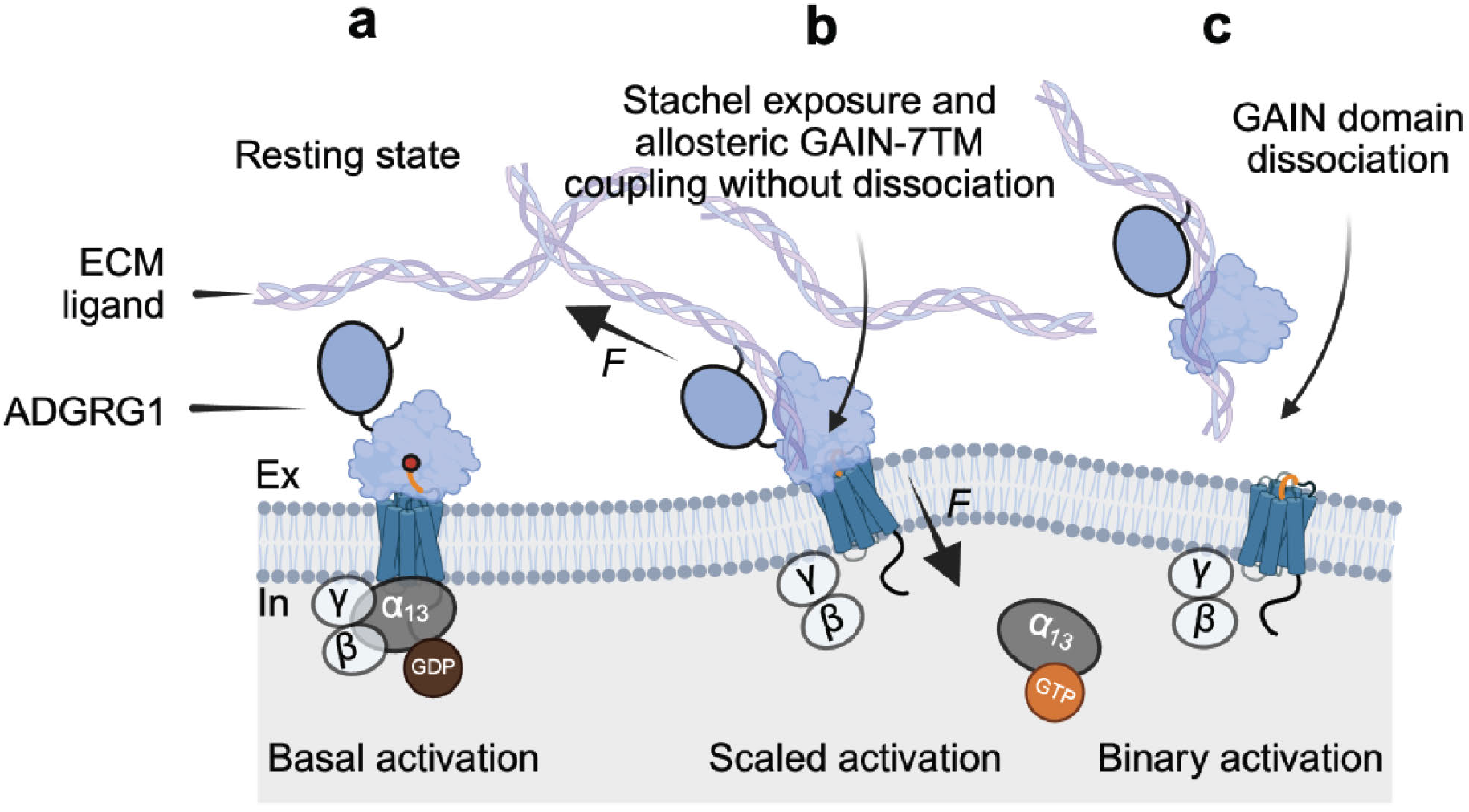
Proposed mechanisms for a dual mode of mechanical-dependent activation of G1. **(a)** G1 undergoes autoproteolytic cleavage at the GPS by its GAIN domain. **(b)** Proposed activation before fragment dissociation. Mechanical stimuli generated by NTF engagement with adhesive ligands and relative movement between the NTF and the membrane-integrated CTF likely alter the interaction between the GAIN domain and the 7TMD. This mechanically induced structural shift may involve partial or complete unfolding of the GAIN subdomain at forces of approximately 5-10 pN, potentially promotes *Stachel* exposure and active allosteric GAIN-7TMD coupling without requiring fragment dissociation, thereby activating G1 in a scaled, potentially reversible, manner. This state is a mechanistic model; its structural basis and signaling reversibility were not directly established here. **(c)** When the applied force exceeds the GAIN domain dissociation threshold (approximately 10-20 pN), the G1 NTF-CTF complex dissociates. The *Stachel* becomes fully exposed and engages the orthosteric binding pocket within the 7TMD, resulting in binary receptor activation. The present experiments do not establish binary signaling behavior at the single-receptor level or a fixed force boundary between the proposed states.

Our live-cell evidence for non-dissociative activation is further supported by receptor biophysics. Complementary force spectroscopy and SMD identified the G1 GAIN domain as a multi-switchable mechanosensor^40^ whose response depends on force direction: some force vectors promote GAIN shedding, whereas physiological pulling geometries stabilize a GAIN-7TMD interface. In the latter state, force reorganizes local hydrogen bonds into a ‘mechanical clamp’ between GAIN loops such as L10 and ECL2, permitting partial *Stachel* exposure and allosteric 7TMD coupling without fragment separation.

Disulfide-locking experiments^40^ further define the required conformational freedom. Internal locking of subdomain A (SS_h1h2) redirects load to subdomain B and favors dissociation and activation, whereas locking subdomain A to L10 (SS_h2l10) prevents shedding and partial unfolding and abolishes activation despite an intact *Stachel*. Thus, an intact TIA/*Stachel* alone is insufficient for signaling; opening of subdomain B/L10 is required to expose it. These findings account for non-dissociative signaling of the G1-H381S mutant: although cleavage-deficient, the receptor remains sufficiently flexible^39^ to undergo force-induced subdomain-B rearrangement.

Mechanistically, our mutagenesis results offer the following interpretations. G1-H381S and G1-T383A may differ in TIA/*Stachel* affinity. T383, the GPS P1’ residue, contributes directly to *Stachel*/peptide docking in the 7TMD^12,20^, accordingly, T383A can retain wild-type basal activity^52,87^ while impairing stimulated signaling. H381S, by contrast, leaves the *Stachel* intact but has been associated with reduced basal signaling^50^ and altered oligomeric dynamics^90^. GAIN-domain unfolding shifts the *Stachel* between encrypted and exposed states; H381S preserves high intrinsic affinity of TIA/*Stachel* and can therefore compensate for reduced exposure compared to CTF-only G1-WT, whereas T383A directly impairs binding^12,20^, with lower affinity. Consistently, the homologous T923G mutation in ADGRL3/LPHN3 impaired the full-length receptor but not the isolated CTF^37^. Alternatively, non-dissociative signalling by cleavage-deficient aGPCRs may not necessarily require *Stachel* engagement with the orthosteric 7TMD pocket, which currently has limited structural support (reviewed in refs. ^81,82^). Rather, an unperturbed *Stachel* could be essential for mediating force-dependent GAIN domain rearrangements that cause allosteric changes of the 7TMD^83^.

The contrasting G1-H381S and G1-T383A phenotypes further argue against a strictly cleavage-dependent model. Autoproteolysis is required for collagen-dependent brain patterning and male fertility in G1-H381S mice^84^ and intraosseous tumor proliferation with G1-T383A^85^, yet uncleaved receptors remain functional in other contexts. For example, the uncleaved G1-H381S receptor supports CNS myelination and cardiac homeostasis in vivo^84^, functions proposed to involve allosteric activation by ligands such as TG2 rather than collagen-like dissociation. Exogenous modulators reinforce this distinction. Synthetic monobodies^51^, small-molecule agonists such as compound 4 (ref.^86^), and activating antibodies^87–89^ stimulate G1-H381S and G1-T383A, demonstrating that G protein activation does not obligatorily require NTF-CTF dissociation. Thus, pharmacological and physiological inputs can access signaling-competent conformations of the intact holoreceptor.

Together with the biophysical findings^40^, these cross-scalar results support a unified model in which the GAIN domain functions as a tunable allosteric mechanosensitive element rather than a static latch, with its flexibility and the vector and magnitude of local force determining whether G1 signals with or without NTF-CTF dissociation: exogenous modulators can shift the 7TMD directly, whereas mechanical stimulation can expose the intact *Stachel* and drive intramolecular coupling without autoproteolysis. The relative contributions of dissociative and non-dissociative activation in full-length WT G1, and their dependence on force magnitude and direction, remain to be determined.

Our novel assays provide a platform to quantify force-response relationships across aGPCRs and to identify allosteric modulators that selectively interfere with mechanotransduction. Future work should quantify how physiological G1 ligands such as collagen III (ref.^46^) and TG2 (ref.^44^) transmit force to the NTF in single-molecule assays. A key unresolved issue is how low-force TIA/*Stachel*-7TMD coupling occurs while the GAIN domain and NTF-CTF complex remain intact; intrinsic GAIN-domain flexibility may permit force-dependent TIA exposure^39–41^.

## MATERIALS AND METHODS

### Cell culture and reagents

COS-7 cells and HFF cells were cultured at 37 ℃ in a 5 % CO2 incubator in high-glucose Dulbecco’s Modified Eagle Medium (DMEM) supplemented with 10 % Fetal Bovine Serum (FBS), 100IU/ml Penicillin/Streptomycin and 1 mM sodium pyruvate (Invitrogen). The cells were tested for mycoplasma contamination and found negative. Human G1 cDNA was cloned into a pcDNA3.1-EGFP vector as described previously^41^. A SpyTag and HaloTag were introduced downstream of the signal peptide sequence in the cDNA to generate the SpyTag-HaloTag-G1-WT-EGFP construct. Mutant SpyTag-HaloTag-G1-H381S/T383A/F385A/(L388A/M389A)-EGFP constructs were created by site-directed mutagenesis using Q5 Site-Directed Mutagenesis Kit (NEB). These mutant constructs were confirmed by Sanger sequencing. pGL4.33[luc2P/SRE/Hygro] were purchased from Promega. HaloTag-miniG13 was a gift from Dr. Nevin, Lambert’s laboratory from Augusta University. For experiments related to the G1 activation, we engineered the G1 constructs by removing the HaloTag to generate SpyTag-G1-WT/H381S/T383A/F385A/(L388A/M389A)-EGFP constructs. G1-EGFP was generated by removing the SpyTag from the SpyTag-G1-EGFP. SpyTag-SNAPTag2-G1-WT/H381S/T383A/F385A/(L388A/M389A)-EGFP constructs were created by replacing the HaloTag to SNAPTag2. COS-7 cells were seeded into a 6-well plate with 60-70 % confluence at day 0 and co-transfected with indicated G1 constructs along with either a HaloTag-miniG13 or a pGL4.33 vector using lipofectamine 3000 (Invitrogen) following the manufacturer’s instructions on day 1. Then the transfected cells were used in the spreading or stretching experiment protocol. A Spy-EGFP construct without the G1 gene was used as an empty vector (EV). Dasatinib (BMS-354825) (100 nM) (Selleckchem) and DHM (10 µM) (Molport) were used in this study.

### Protein expression and purification

pET28a(+) plasmids containing Biotin-SpyCacher, Biotin-TG2(D3D4), and Biotin-TG2(D3D4)-mScarlet3 domain were transformed in *E.coli* BL21(DE3) cells with biotin protein ligase (BirA) and purified using a His-tag affinity column following the previous protocol^91^. Colonies transformed with the respective plasmids were first grown overnight in 10 mL LB medium containing 100 µg/mL kanamycin at 37 °C. The overnight cultures were then inoculated into 1 L of fresh LB medium with ampicillin and incubated at 37 °C for 4-6 hours until the optical density at 600 nm (OD600) reached ∼0.6. Protein expression was induced by adding 0.4 mM IPTG and 0.5 mM biotin, followed by overnight incubation at 20 °C. Biotinylation was catalyzed by BirA, enabling site-specific conjugation to the AviTag. Bacterial cells were harvested by centrifugation at 6000 g and pellets were resuspended in lysis buffer (50 mM Tris, 300 mM NaCl, pH 7.4) containing 1 mM PMSF, and lysed using a French press.

Lysates were centrifuged at 40,000 g for 30 minutes, and the supernatant was incubated with Co^2+^-NTA resin (Thermo Scientific) for 1 hour. After washing with buffer 50 mM Tris, 300 mM NaCl, 10 mM imidazole, pH 7.4), proteins were eluted using buffer containing 200 mM imidazole. The eluted proteins were further purified by size-exclusion chromatography (Superdex 200, AKTA Pure, GE Healthcare, MA). Protein-containing fractions were analyzed by SDS-PAGE, dialyzed into PBS, aliquoted with 10% (v/v) glycerol, flash-frozen in liquid nitrogen, and stored at -80 °C. Protein concentration was determined by BCA assays (Thermo Scientific).

Codon-optimized cDNA encoding the human GPR56 fragment (residues 173-394) was synthesized and cloned into a customized pLEXm vector containing an N-terminal signal peptide and a C-terminal His-tag. An AviTag and a SpyTag were introduced at the N- and C-termini, respectively. The H381S and T383A point mutants were generated by site-directed mutagenesis and cloned into the same vector.

HEK293F GnTI− cells were cultured in FreeStyle 293 Expression Medium at 37 °C, 5% CO₂, and 95% humidity on an orbital shaker with a 50-mm amplitude. At a density of 1–2×10^6^ cells/mL, the cells were transiently transfected using DNA and polyethylenimine at a mass ratio of 1:3. After 3-5 days, the culture supernatants were collected and incubated with Ni-NTA beads (Smart-Lifesciences, China) at 4 °C for approximately 2 h. The beads were washed with 20 mM HEPES, pH 7.5, 500 mM NaCl, and 30 mM imidazole, and the bound proteins were eluted using the same buffer containing 300 mM imidazole. The proteins were further purified by size-exclusion chromatography using a Superdex 200 Increase column (Cytiva) equilibrated with 20 mM HEPES, pH 7.5, and 150 mM NaCl. Purified proteins were concentrated, biotinylated using a BioA kit (Avidity), and evaluated by Western blotting. Successfully biotinylated samples were used for magnetic-tweezers experiments. All mutant proteins were expressed and purified using the same procedures.

### Cell migration and stretching

5×10^3^ transfected single COS-7 or HFF cells were seeded on the glass bottom dishes (ibidi), or spin-coated with a layer of rigid PDMS (2MPa) as described previously^92^, and coated with Fibronectin (Roche) (10 µg/ml), Fibronectin together with TG2 (D3D4) domain (20 µg/ml), or Fibronectin together with SpyCatcher (20 µg/ml) in PBS for 1 hour at room temperature. The cells were incubated with a Janelia Fluor -646 (JF646) HaloTag Ligands (Promega) for 1 hour before cell spreading. After 3 hours of cell spreading, the process of cell migration on PDMS surfaces was performed on W1 Spinning disc confocal microscopy for visualizing the real-time GAIN domain dissociation (1 hour imaging) and on widefield microscopy EZ (Olympus IX81) for calculating the migration speed (24 hours). After overnight cell migration, the cells were fixed for immunofluorescence. The co-localization of G1-EGFP with the NTF of G1 were performed on W1 Spinning disc confocal microscopy (PDMS surfaces) or on the total internal reflection fluorescence microscopy (TIRFM) (glass surfaces). The dynamic recruitment of miniG13 during cell migration (after 3 hours of cell spreading) on the glass surfaces were monitored with a TIRFM. Imaging was done on a Nikon TIRF system with 488 and 640 laser lines. For cell stretching, 5×10^3^ transfected COS-7 cells were seeded on the cell stretching system (STREX) coated with Fibronectin or Fibronectin together with SpyCatcher. Cells were then stretched to 16% strain for 1h and were fixed for immunofluorescence.

### Micropattern fabrication and imaging

The coverslips (25 mm in diameter, Paul Marienfeld) were initially immersed in a detergent bath (50% Decon 90, Decon Laboratories Limited) to eliminate surface impurities and underwent a 30-minute sonication to ensure thorough cleaning. The samples were subsequently rinsed multiple times with deionized (DI) water, ethanol, and acetone, then dried using a nitrogen stream. To generate surface hydroxyl groups, the dried glasses were subjected to oxygen plasma treatment (8 %, 50 mW, Tergeo plasma cleaner) for 10 minutes. Next, the substrates were submerged in an ethanol solution containing 1 % (v/v) APTES (3-aminopropyltriethoxysilane) at room temperature for 30 minutes, rinsed again with ethanol, dried under nitrogen flow, and heated at 90 °C for 1 hour to achieve amino group functionalization. Then the coverslips were coated with 1 % (v/v) glutaraldehyde solution for 30 minutes, washes five times with 1× PBS, and dried using a nitrogen stream.

Master molds containing the intended micropattern design (12 µm circles separated by 10 µm gaps) were fabricated on silicon wafers using SU8-3050 photoresist and conventional photolithography. Polydimethylsiloxane (PDMS) stamps were then cast from these molds and employed for microcontact printing. The PDMS stamps were incubated with NeutrAvidin (0.1 mg/ml) for 30 minutes and subsequently brought into contact with pre-treated coverslips to transfer the pattern. After one hour of contact, the stamps were gently removed to preserve the printed features. The patterned coverslips were coated with fibronectin (10 µg/ml) for 30 minutes, followed by overnight blocking with 3% BSA in PBS. Finally, specific biotinylated proteins of interest (e.g., Biotin-SpyCatcher or Biotin-TG2 (D3D4)) were applied at a concentration of 50 µg/ml to generate functionalized micropatterns.

COS-7 cells transfected with both HaloTag-miniG13 and SpyTag-SNAPTag2-G1-WT/H381S/T383A/F385A/(L388A/M389A)-EGFP were incubated with SNAP-Cell TMR-Star (New England Biolabs) for 30 min, followed by three washes with culture medium. Cells were then incubated with Janelia Fluor 646 (JF646) HaloTag ligand (Promega) for 1 hour before cell seeding. 3×10^3^ transfected single COS-7 cells were seeded on the micro-patterned surface. After 3 hours of cell spreading, real-time monitoring of GAIN domain dissociation and dynamic miniG13 recruitment during cell migration was performed using total internal reflection fluorescence microscopy (TIRFM). Imaging was conducted on a Nikon TIRF system equipped with 488, 555, and 640 nm laser lines.

### Magnetic tweezer-based single-molecule force approach

Single-molecule manipulation experiments were performed using a custom-built magnetic-tweezers setup according to previously described procedures^41^. Force was calibrated individually for each tethered bead, with an estimated relative uncertainty of approximately 10%. The applied force, *f*, was determined from the distance, (*d*), between the permanent magnets and the paramagnetic bead. The force-distance relationship was calibrated using 16-μm λ-DNA. For the magnet pair used in this study, the relationship was described by:

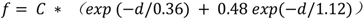

Here, (*d*) is expressed in millimeters, and (*C*) is a bead-specific calibration constant. For M-270 paramagnetic beads, (*C*) ranged from 140 to 180.

### Immunofluorescence

After undergoing cell spreading or stretching, COS-7 or HFF cells were fixed with a pre-warmed solution of 4 % paraformaldehyde in PBS at 37°C for 15 minutes. This was followed by permeabilization with 0.2 % Triton X-100 in TBS for 30 minutes at room temperature to allow for antibody penetration. To minimize non-specific binding, the samples were then treated with a blocking solution of 1% Bovine Serum Albumin (BSA) in TBS for 1 hour. The samples were then incubated with primary antibodies (Anti-G1 Antibody (G-6) N-terminal (Santa Cruz, sc-390192), diluted 1:200, integrin α5 (BD Pharmingen, Purified NA/LE (CD49e), 555614), diluted 1:200) overnight at 4°C, followed by incubation with secondary antibodies and Alexa Fluo Plus 405 Phalloindin (Invitrogen) for 1 hour at room temperature. The co-localization of G1-EGFP with the NTF of G1 was analyzed by calculating Pearson’s correlation coefficient between the two channels.

### Western blot

Confluent cells were lysed in a RIPA buffer (Sigma) and a protease inhibitor (Roche) for 20 minutes at 4 °C. The amount of proteins extracted was determined using the BCA assay (Bio-Rad). The extracted proteins (20 μg) were separated by 4-20% SDS polyacrylamide gel electrophoresis (PAGE) and transferred to nitrocellulose membranes (Bio-Rad) using an electro-transfer process for 7 minutes (Trans-Blot Turbo system). The membranes were blocked with a solution of 5 % Bovine Serum Albumin (BSA) in TBS 0.05 % Tween 20 to prevent non-specific binding. The membranes were then incubated with primary antibodies (Anti-G1 Antibody (G-6) N-terminal (Santa Cruz, sc-390192), diluted 1:1000) overnight at 4 °C, followed by an incubation with secondary HRP antibodies (diluted 1:2000) for 1 hour at room temperature. The results were visualized using the ChemiDoc chemiluminescence detection system (Bio-Rad) and quantified using Image Lab. GAPDH was used as a reference protein to control for variations in protein loading.

### ELISA

A total of 2.5×10^4^ COS-7 cells were seeded per well in a 96-well white plate (Costar, #3610) pre-coated with 0.1 mg/mL poly-L-lysine (PLL; Sigma). Twenty-four hours after seeding, cells were transfected as previously described with 0.1 μg of the indicated SpyTag-G1^x^-EGFP construct and 25 ng of pGL4.33[luc2P/SRE/Hygro] (Promega). After an additional 24 hours, cells were washed with PBS and fixed with cold 4 % paraformaldehyde (PFA) for 10 min at 4 °C. Following fixation, cells were washed again with PBS to remove residual PFA. Cells were then subjected to three sequential 1-hour incubations: (1) blocking with 2 % BSA in PBS; (2) incubation with primary anti-G1 antibody (G-6, N-terminal; Santa Cruz Biotechnology, sc-390192) at 0.05 μg/mL (1:4000 dilution) in 2% BSA-PBS; and (3) incubation with HRP-conjugated anti-mouse secondary antibody (Cell Signaling Technology) at a 1:5000 dilution in 2 % BSA-PBS. Cells were thoroughly washed with PBS following each antibody incubation step. Subsequently, cells were incubated with Clarity Western ECL Substrate (Bio-Rad) at room temperature for 10 min, and luminescence was measured using a GloMax® Discover microplate reader (Promega). All assays were performed in triplicate. Luminescence signals were normalized to the background signal obtained from cells transfected with Spy-EGFP, which served as the empty vector (EV) control.

### Dual-Glo SRE-Luciferase Assay

The co-transfected COS-7 cells in the 6-well plate were subjected to 16 hours of serum starvation. These cells (7.5×10^4^) were then seeded on glass bottom dishes that were coated with either Fibronectin, Fibronectin together with TG2 (D3D4), or Fibronectin together with SpyCatcher to promote cell spreading. After 3 hours of cell spreading, the cells were lysed and analyzed using the Dual-Glo luciferase assay system (E2920, Promega), according to the manufacturer’s protocol. For the stretched cell experiment, co-transfected COS-7 cells were seeded on a cell stretching system (STREX) that was coated with either FN or FN+Spy and subjected to 16 hours of serum starvation. The cells were then stretched at 16 % strain for 1 hour, followed by 2 hours of recovery, and lysed and analyzed using the Dual-Glo luciferase assay system. The luminescence readings were taken using a Glomax Discover plate reader (Promega). The firefly luciferase data was normalized to the Renilla luciferase signal and expressed as the fold increase over the signal obtained from cells that were transfected with Spy-EGFP as an Empty Vector (EV) underwent the same spreading or stretching.

### Statistical analyses

To capture the dynamic changes in G1 and miniG13 colocalization, several Spy adhesions sites (Fig. 4c-d) or TG2 micropatterned areas (Fig. 5e-f) from each cell were analyzed, and the average fluorescence signal intensities were normalized to the corresponding background average intensities outside Spy adhesion sites or TG2 micropatterned areas. The obtained G1 enrichment ratio is noted as I_EGFP_, and the miniG13 enrichment ratio is I_miniG13_, respectively, with I_EGFP,0_ and I_miniG13,0_ indicating the normalized intensities at the start of a 1 h imaging time interval, during which the dynamic changes in I_EGFP_/I_EGFP,0_ (green) and I_miniG13_/I_miniG13,0_ (magenta) were plotted at a rate of 2 frame/min. Each data point represents the average of > 30 Spy adhesion sites (n = 5 sites/cell; n > 6 cells) or > 15 TG2 micropatterned areas.

To measure the dynamic changes of NTF/CTF ratio, several TG2 (D3D4) (Fig. 5c-d) or SpyCatcher (Extended Data Fig. 4a, c) coated micropatterned areas from each cell were analyzed, and the average fluorescence signal intensities were normalized to the corresponding background average intensities outside TG2 or Spy micropatterned areas. The obtained G1-NTF signal is noted as I_SNAP2_, and the G1-CTF signal is I_EGFP_, respectively, with I_SNAP2,t_ and I_EGFP,t_ indicating the normalized intensities at the indicated time of a 1 h imaging time interval, during which the dynamic changes of NTF/CTF ratio is calculated as I_SNAP2,t_/I_EGFP,t_, plotted at a rate of 2 frame/min. Each data point represents the average of > 10 TG2 or > 8 Spy micropatterned areas.

To ensure accuracy and reproducibility, measurements were performed two to three times in triplicate. Prism (GraphPad Software) was used for data analysis and graph plotting. The data was visualized using Adobe Illustrator. The illustrations were drawn with the help of Biorender. To determine statistical significance, an ANOVA test and an unpaired Welch’s t-test were conducted. The results were considered statistically significant if P is less than 0.05.

### G1 GAIN domain partial and full unfolding calculation

The G1 GAIN domain comprises 222 amino acids, corresponding to a theoretical contour length of approximately 84.4 nm, based on a contour-length increment of 0.38 nm per residue. It can be divided into subdomain A, comprising 49 residues with a theoretical contour length of approximately 18.6 nm, and subdomain B, comprising 173 residues with a theoretical contour length of approximately 65.7 nm. Subdomain B contains the 12-residue TIA, which forms its C-terminal β-strand. Excluding the *Stachel* sequence, subdomain B comprises 161 residues, corresponding to a contour length of approximately 61.2 nm.

The experimentally released contour length of the G1 GAIN domain was determined by summing the contour-length increments associated with individual unfolding events. For example, the H381S mutant exhibited single-step unfolding transitions with an unfolding force of 7.5 ± 3.9 pN and a step size of 40.1 ± 8.4 nm. The corresponding released contour length was 76.9 ± 5.7 nm, consistent with complete unfolding of the G1 GAIN domain. Using the partial unfolding force (5.8 ± 3.0 pN) and unfolding step size (21.5 ± 11.0 nm) of H381S mutant, we calculated a released contour length of 45.2 ± 23.2 nm. This value exceeds the contour length of GAIN subdomain A (18.6 nm) but is smaller than that of subdomain B (65.7 nm). The mean released contour length exceeds the theoretical contour length of subdomain A alone and is consistent with unfolding involving subdomain B. However, contour length alone does not identify the unfolded segments or establish *Stachel* exposure.

#### Protein sequences

Biotin(AviTag)-SpyCacher, Biotin(AviTag)-TG2(D3D4), and Biotin(AviTag)-TG2(D3D4)-mScarlet3 domain, Biotin(AviTag)-G1-GAIN-SpyTag sequence were used in this study.

Biotin(AviTag)-SpyCacher

MGSSHHHHHHSSGLVPRGSHMASMTGGQQMGRDPNSSSVDKLMCGSSGLNDIFEAQKIEWHESSGENLYFQSGAMVDTLSGLSSEQGQSGDMTIEEDSATHIKFSKRDEDGKELAGATMELRDSSGKTISTWISDGQVKDFYLYPGKYTFVETAAPDGYEVATAITFTVNEQGQVTVNGKATKGD AHI*

Biotin(AviTag)-TG2(D3D4)

MGSSHHHHHHSSGLVPRGSHMASMTGGQQMGRDPNSSSVDKLMCGSSGLNDIFEAQKIEWHESSGENLYFQSGSGSSGSGSGGSSGSGGSSGSGSSGSGSSGSSGSSGSGSGGSGSSGSGGGLAEKEETGMAMRIRVGQSMNMGSDFDVFAHITNNTAEEYVCRLLLCARTVSYNGILGPECGTKYLLNLNLEPFSEKSVPLCILYEKYRDCLTESNLIKVRALLVEPVINSYLLAERDLYLENPEIKIRILGEPKQKRKLVAEVSLQNPLPVALEGCTFTVEGAGLTEEQKTVEIPDPVEAGEEVKVRMDLLPLHMGLHKLVVNFESDKLKAVKGFRNVIIGPA*

Biotin(AviTag)-TG2(D3D4)-mScarlet3

MGSSHHHHHHSSGLVPRGSHMASMTGGQQMGRDPNSSSVDKLMCGSSGLNDIFEAQKIEWHESSGENLYFQSGSGSSGSGSGGSSGSGGSSGSGSSGSGSSGSSGSSGSGSGGSGSSGSGGGLAE KEETGMAMRIRVGQSMNMGSDFDVFAHITNNTAEEYVCRLLLCARTVSYNGILGPECGTKYLLNLNLEPFSEKSVPLCILYEKYRDCLTESNLIKVRALLVEPVINSYLLAERDLYLENPEIKIRIL GEPKQKRKLVAEVSLQNPLPVALEGCTFTVEGAGLTEEQKTVEIPDPVEAGEEVKVRMDLLPLHMGLHKLVVNFESDKLKAVKGFRNVIIGPAGSIGSMDSTEAVIKEFMRFKVHMEGSMNGHEFEIEGEGEGRPYEGTQTAKLRVTKGGPLPFSWDILSPQFMYGSRAFTKHPADIPDYWKQSFPEGFKWERVMNFEDGGAVSVAQDTSLEDGTLIYKVKLRGTNFPPDGPVMQKKTMGWEASTERLYPEDVVLKGDIKMALRLKDGGRYLADFKTTYRAKKPVQMPGAFNIDRKLDITSHNEDYTVVEQYERSVARHSTEFGGGS*

Biotin(AviTag)-hG1-GAIN (173^rd^-394^th^)-SpyTag

LNDIFEAQKIEWHEGGGSGSVDMSELKRDLQLLSQFLKHPQKASRRPSAAPASQQLQSLESKLTSVRFMGDMVSFEEDRINATVWKLQPTAGLQDLHIHSRQEEEQSEIMEYSVLLPRTLFQRTKGRSGEAEKRLLLVDFSSQALFQDKNSSQVLGEKVLGIVVQNTKVANLTEPVVLTFQHQLQPKNVTLQCVFWVEDPTLSSPGHWSSAGCETVRRETQTSCFCNHLTYFAVLMVSSVEGGGSGAHIVMVDAYKPTKHHHHHHA*

H381S: H→S; T383A: T→A

## DATA AVAILABILITY

The raw Western blot data are available in figshare at DOI: https://doi.org/10.6084/m9.figshare.33721534. All other data are available upon request to the corresponding authors.

## ACKNOWLEDGEMENTS

This work was supported through grants of the Deutsche Forschungsgemeinschaft (SFB1423, project number 421152132, subproject A06; project number 542631574) to T.L., Singapore Ministry of Education Academic Research Fund Tier 3 (MOE Grant No: MOET32021-0003) and Tier 2 (MOE Grant Nos: MOE-T2EP50123-0008), National Research Foundation (NRF) Singapore, Mechanobiology Institute under its MID-SIZED GRANT (MSG) (NRF-MSG-2023-0001) to J.Y. We thank Dr. Nevin, Lambert (Augusta University) for sharing miniG plasmids and Dr. Xianhua Piao (University of California San Francisco) for sharing the G1 plasmids, and MBI wet lab and imaging core facilities.

## AUTHOR CONTRIBUTIONS

C.F., T.L., and J.Y. designed research

C.F., and C.Q. performed the cell migration and deformation experiments

C.F., and L.Z. performed the magnetic tweezer experiments

Y.W. expressed and purified the isolated G1 GAIN domains

C.F., C.Q., L.Z., and Y.Z. analyzed the data

T.L., and J.Y. co-supervised the research

C.F., G.S., T.L., and J.Y. wrote the paper

## ADDITIONAL INFORMATION

## Supplementary information

The online version contains supplementary material.

### Correspondence and requests for materials

Correspondence and requests for materials should be addressed to Yan Jie and Tobias Langenhan.

## EXTENDED DATA FIGURE LEGENDS

**Extended Data Figure 1.**
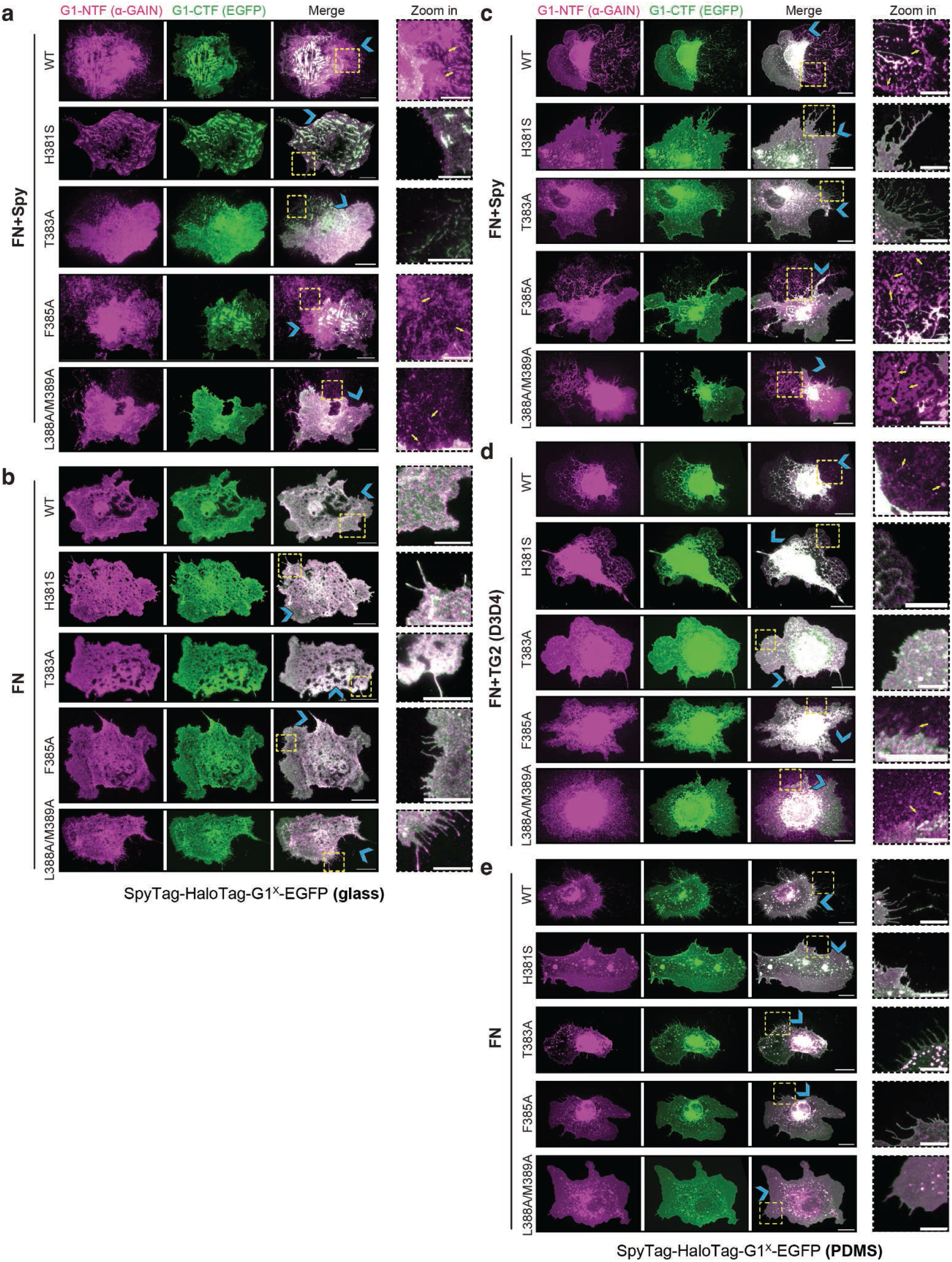
G1 GAIN domain dissociation probed with ‘migration force assay’. (a-e) Immunostaining of G1-NTF (α-GAIN, magenta) and fluorescence imaging of G1-CTF (EGFP, green) in COS-7 cells expressing the indicated SpyTag-HaloTag-G1^x^-EGFP variants and spreading on FN+Spy **(a)** and FN **(b)** coated glass surfaces, FN+Spy **(c)**, FN+TG2 (D3D4) **(d)**, and FN **(e)** coated PDMS surfaces. Blue chevrons indicate migration direction. Yellow arrows highlight dissociated NTFs (zoom-in). Scale bar: 20 µm; zoom-in: 10 µm.

**Extended Data Figure 2.**
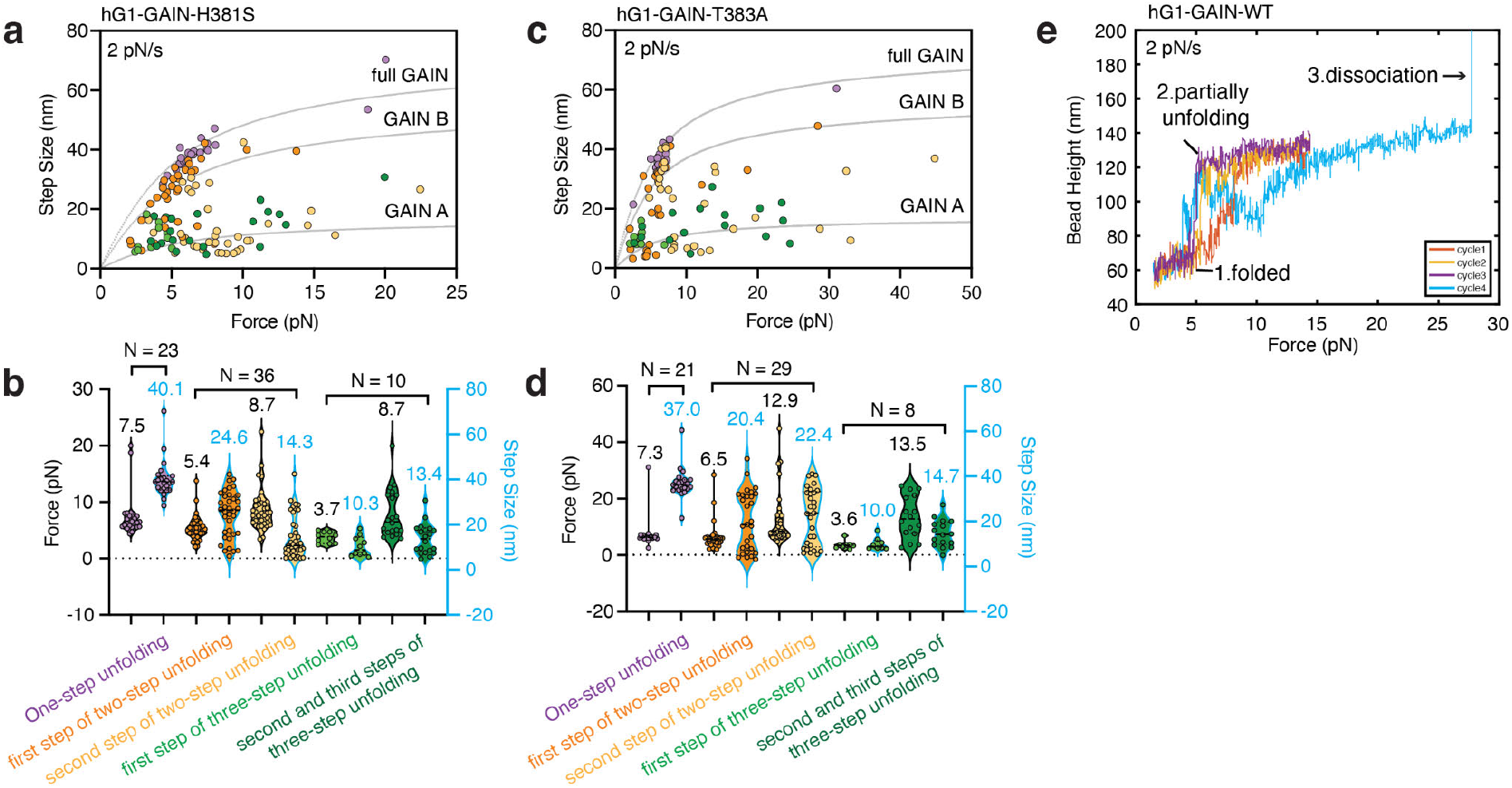
Force induced GAIN domain dissociation, partial or full unfolding in G1 variants. (a, **c)** Scatter plots of unfolding force versus step size for the H381S (**a**) and T383A (**c**) GAIN domains. Orange and light-green circles indicate the first unfolding transitions, whereas yellow and dark-green circles indicate the second transitions, which are occasionally separated into two substeps. Purple circles represent combined multistep unfolding events (N=23). **(b, d)** Violin plots summarizing the distributions of unfolding forces and step sizes for H381S (**b**) and T383A (**d**). Mean values are indicated, and N denotes the number of force-loading cycles. Data were obtained from eight H381S tethers and five T383A tethers across at least three independent experiments. **(e)** Four representative force-height traces recorded for the WT G1 GAIN domain during repeated loading cycles at 2 pN/s. The force did not exceed 15 pN during cycles 1-3, allowing unfolding without dissociation. During cycle 4, the force was increased beyond this range, resulting in NTF-CTF dissociation.

**Extended Data Figure 3.**
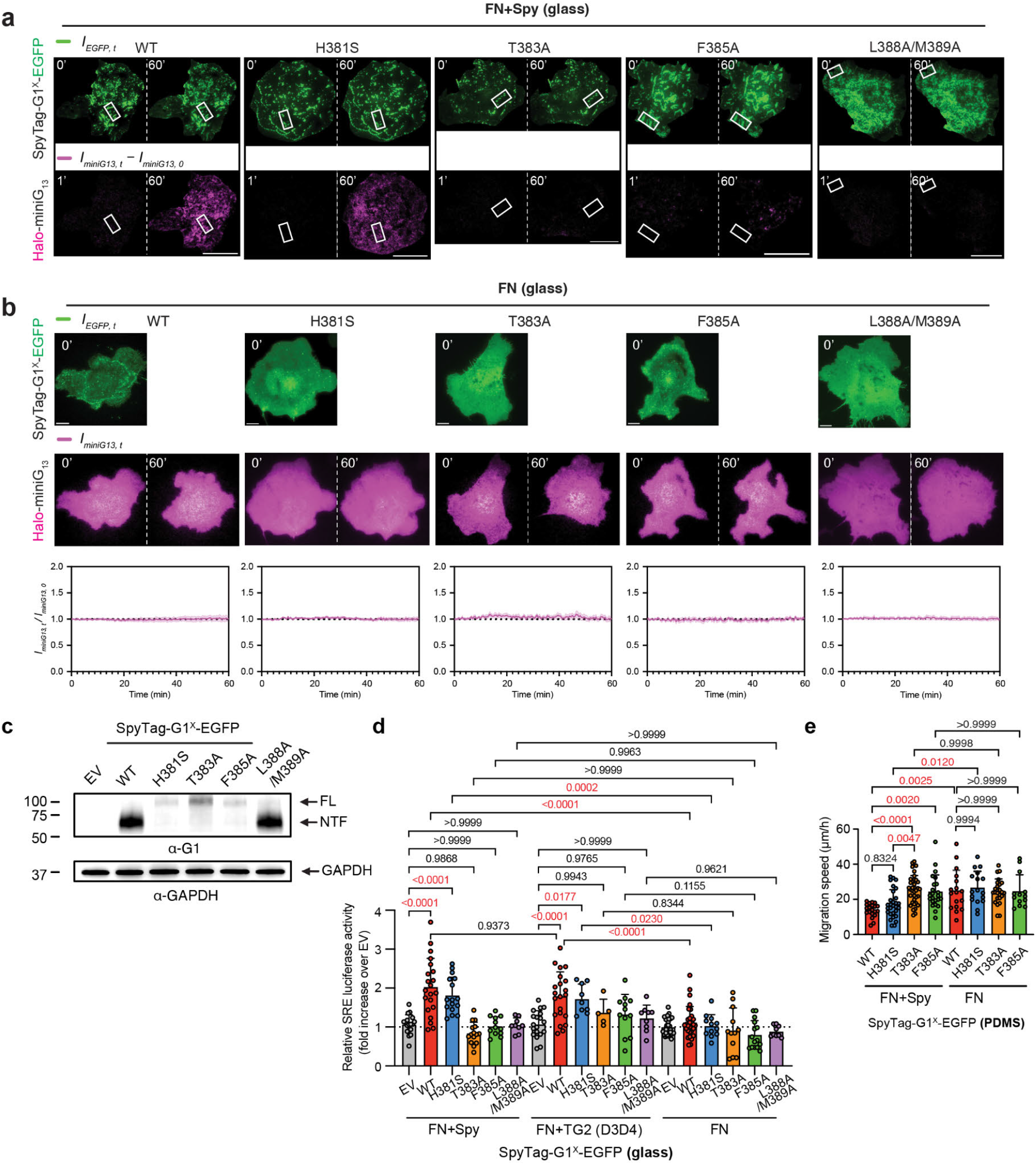
Dynamic mechanical activation of G1 during cell migration. **(a)** TIRFM images of COS-7 cells expressing SpyTag-G1^x^-EGFP variants (green) migrating on FN+Spy-coated glass surfaces. HaloTag-miniG13 (magenta) recruitment is recorded. The net change in HaloTag-miniG13 fluorescence between the indicated time points was shown. Scale bar: 10 µm. **(b)** TIRFM images of cells expressing SpyTag-G1^x^-EGFP variants (green) migrating on FN surfaces at indicated time points, showing no HaloTag-miniG13 (magenta) recruitment over 60 minutes. Scale bar: 10 µm. Dynamic changes in I_miniG13,t_/I_miniG13,0_ (magenta) are shown; n ≥ 5 cells per group. **(c)** Western blot analysis of COS-7 cells expressing SpyTag-G1 variants (G1^x^ corresponds to GPS mutations). **(d)** Relative luciferase activity for SpyTag-G1^x^-EGFP variants on FN+Spy, FN+TG2 (D3D4 domains) or FN-coated glass surfaces during migration. **(e)**. Migration speed of COS-7 cells expressing SpyTag-G1^x^-EGFP variants on FN or FN+Spy surfaces. Data (n ≥ 13 cells) are shown as scatter plots with mean ± SD. Statistical analysis: one-way ANOVA with Tukey’s test (P values shown).

**Extended Data Figure 4.**
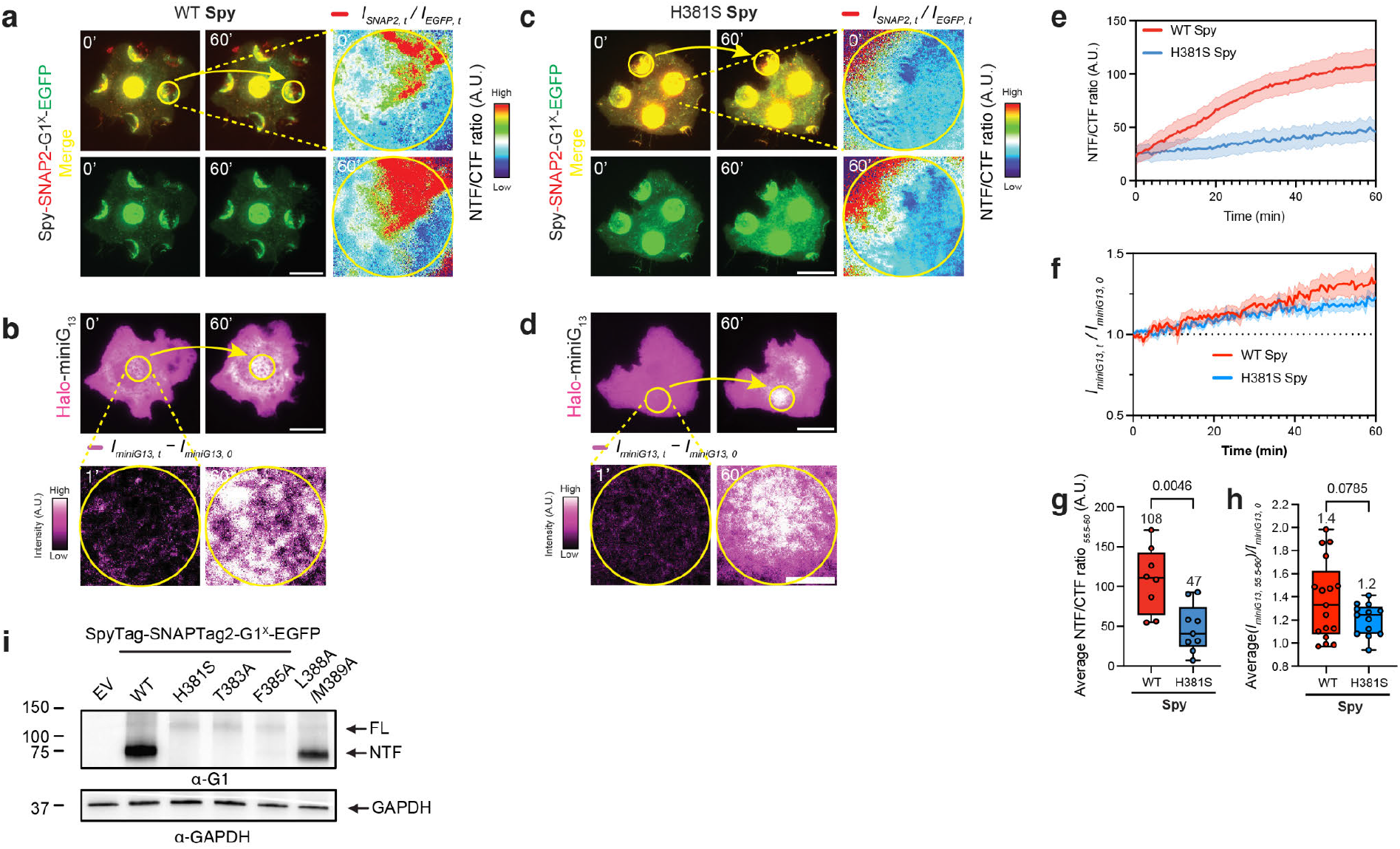
G1 dissociation occurs on WT but not H381S and both show mechanical activation on Spy-coated micropatterns. (a, **c)** TIRFM images of COS-7 cells expressing SpyTag-SNAPTag2-G1^x^-EGFP variants (yellow: Merge of SNAPTag2, red, and EGFP, green) migrating on Spy-coated micropatterns. Scale bar: 20 µm. The yellow circle (12 µm diameter) indicates the zoomed-in region showing G1 dissociation. The NTF/CTF fluorescence ratio reflects the extent of G1 dissociation. Scale bar: 5 µm. G1 dissociation occurs in WT but not in H381S. **(b, d)** TIRFM images of COS-7 cells expressing SpyTag-SNAPTag2-G1^x^-EGFP variants migrating on Spy-coated micropatterns. Scale bar: 20 µm. HaloTag-miniG13 (magenta) is recruited to both WT G1 and the H381S mutant. The yellow circle (12 µm diameter) indicates the zoomed-in region showing miniG13 recruitment at Spy-coated micropatterns. The miniG13 intensity represents the net increase during the 60-minute interval. Scale bar: 5 µm. **(e)** Time traces of NTF/CTF fluorescence intensity showing G1 adhesion dynamics at SpyCatcher-coated micropatterns over 60 minutes. **(f)** Dynamic changes in miniG13 intensity (I_miniG13,t_/I_miniG13,0_, magenta) at SpyCatcher-coated micropatterns over 60 minutes. **(g)** Average NTF/CTF ratio and **(h)** I_miniG13,t_/I_miniG13,0_ values from the final 5 minutes of imaging in **(e-f)**. Data (n ≥ 10 replicates) are shown as scatter plots with mean ± SD. Statistical analysis: Welch’s t-test (P values shown). **(i)** Western blot analysis of COS-7 cells expressing SpyTag-SNAPTag2-G1 variants (G1^X^ corresponds to GPS mutations).

**Extended Data Figure 5.**
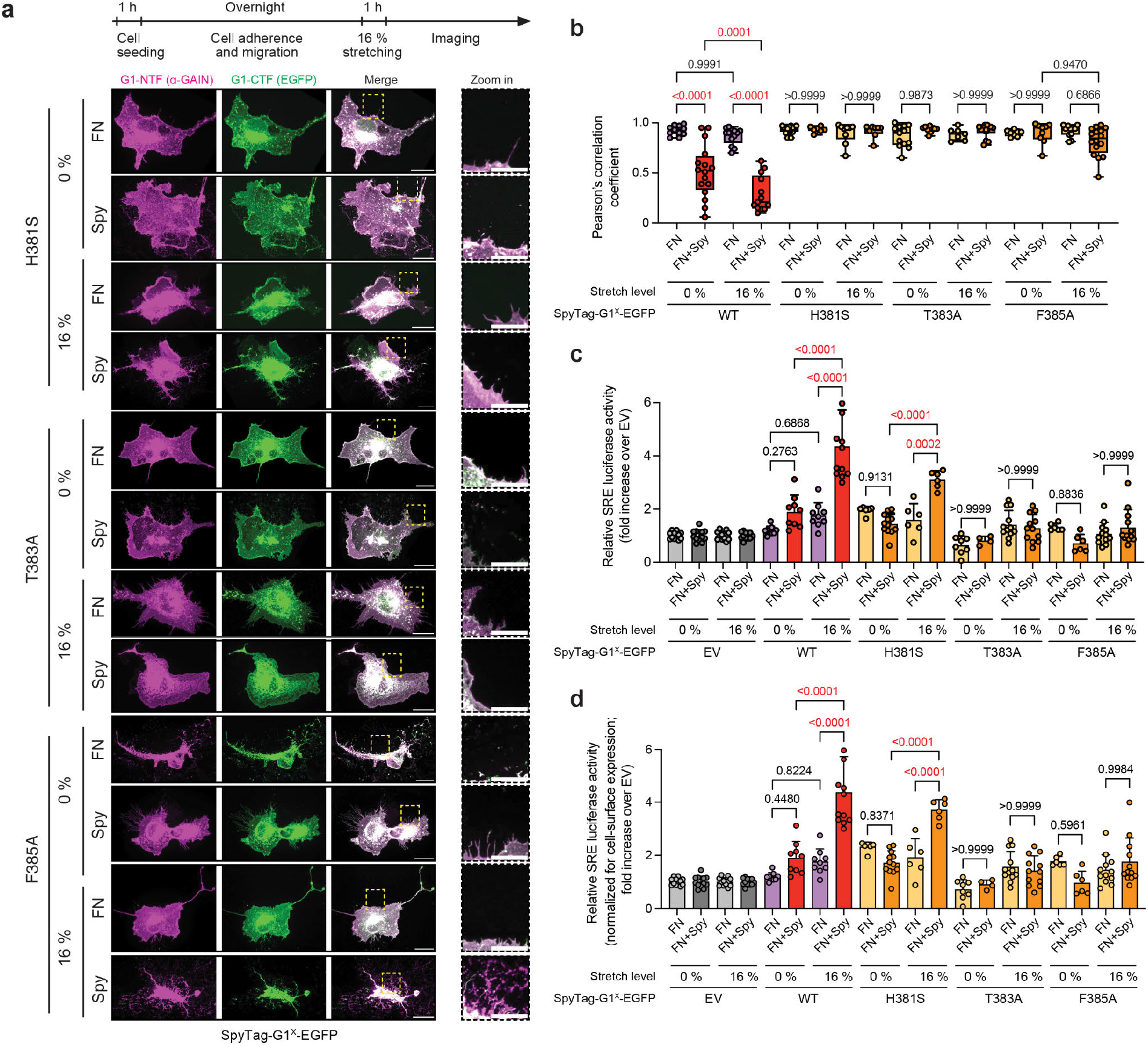
G1 dissociation and activation with ‘deformation force assays’. **(a)** Immunostaining of COS-7 cells expressing SpyTag-G1^x^-EGFP variants on FN or FN+Spy surfaces, either unstretched or stretched to 16 %. NTF (α-GAIN, magenta) and CTF (EGFP, green) are visualized. Scale bars: 20 µm (main) and 10 µm (zoom). **(b)** Correlation between NTF (α-GAIN) and CTF (EGFP) signals from **(a)**, quantified with Pearson’s correlation coefficient. Data (>10 biological replicates from 3-independent experiments) are shown as box-and-whisker plots (all data points plotted; median: horizontal line; boxes: 25th-75th percentiles; whiskers: min and max). Statistical analysis: one-way ANOVA with Tukey’s multiple comparisons test. P-values indicated. For reference, the WT dataset from Fig. 6c is displayed in this extended dataset. **(c)** G1 activation under deformation forces assessed via SRE-luciferase reporter assay for conditions in **(a)**. **(d)** Mutant constructs normalized for cell-surface expression from Fig. 4i. For reference, the full EV and WT datasets from Fig. 6d and the 16 % data for all receptor versions in Fig. 6e are displayed in this extended dataset. EV served as a control. Data (n > 6 replicates, 3-independent experiments) are presented as scatter plots with bars (mean ± SD). Statistical analysis: one-way ANOVA with Tukey’s test. P-values indicated.

## SUPPLEMENTARY MATERIAL

**Supplementary Figure 1.**
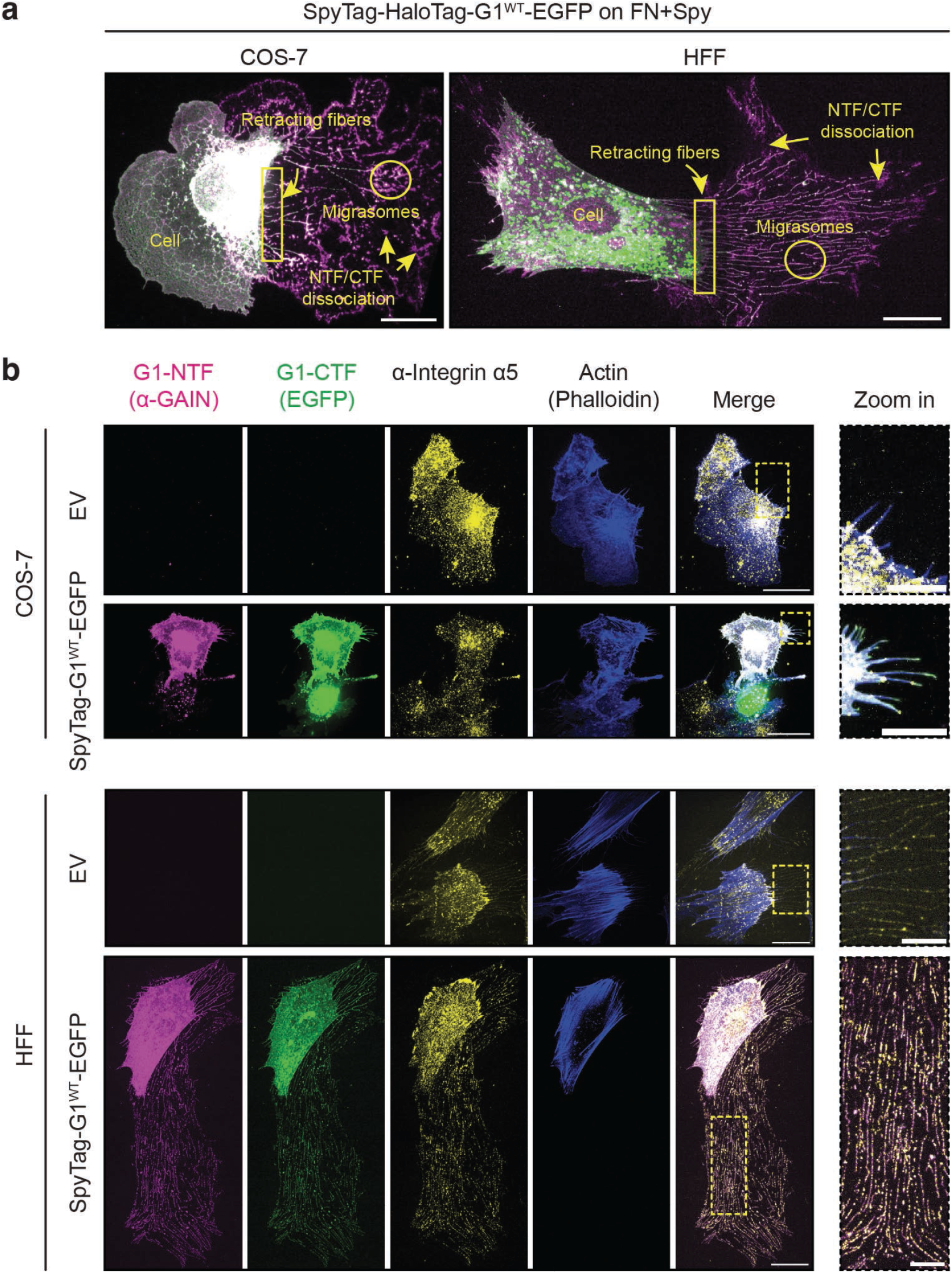
Formation of retracting fibers, migrasomes and occurrence of G1 dissociation during cell migration. **(a)** Immunostaining of COS-7 and HFF cells expressing SpyTag-HaloTag-G1^WT^-EGFP on FN+Spy surfaces, showing the cell body, retracting fibers, migrasomes, and NTF/CTF dissociation. Scale bar: 20 µm. **(b)** Immunostaining of COS-7 and HFF cells expressing pCDNA3.1 (EV) or SpyTag-G1-EGFP on FN surfaces, highlighting NTF (α-GAIN), CTF (EGFP), integrin α5, and actin (Phalloidin). Scale bars: 30 µm (overview) and 10 µm (zoom-in).

**Supplementary Figure 2.**
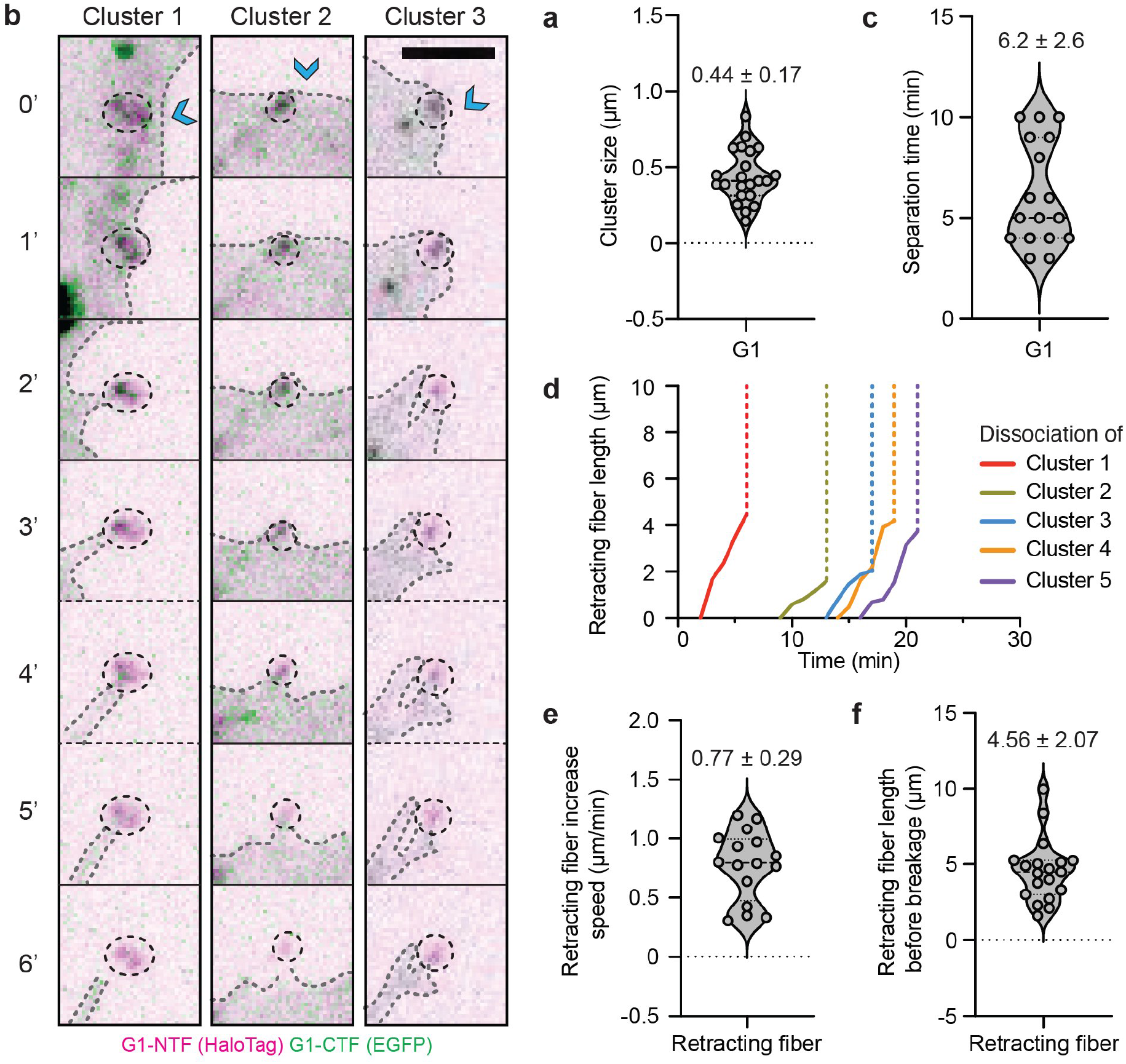
G1 dissociation occurs at retracting fibers within minutes after formation. **(a)** Cluster size quantification of SpyTag-HaloTag-G1-EGFP. **(b)** Video frames showing NTF-CTF dissociation of SpyTag-HaloTag-G1-EGFP clusters over 1’ intervals. Blue chevrons indicate cell migration direction. Scale bars: 2.5 µm. **(c)** Estimated NTF-CTF separation time during cell migration on FN+Spy surfaces (violin plot). **(d)** Representative traces of retracting fiber length increase over time before breakage. **(e)** Local retraction speed and **(f)** retracting fiber length during cell migration on FN+Spy surfaces (violin plots).

**Supplementary Figure 3.**
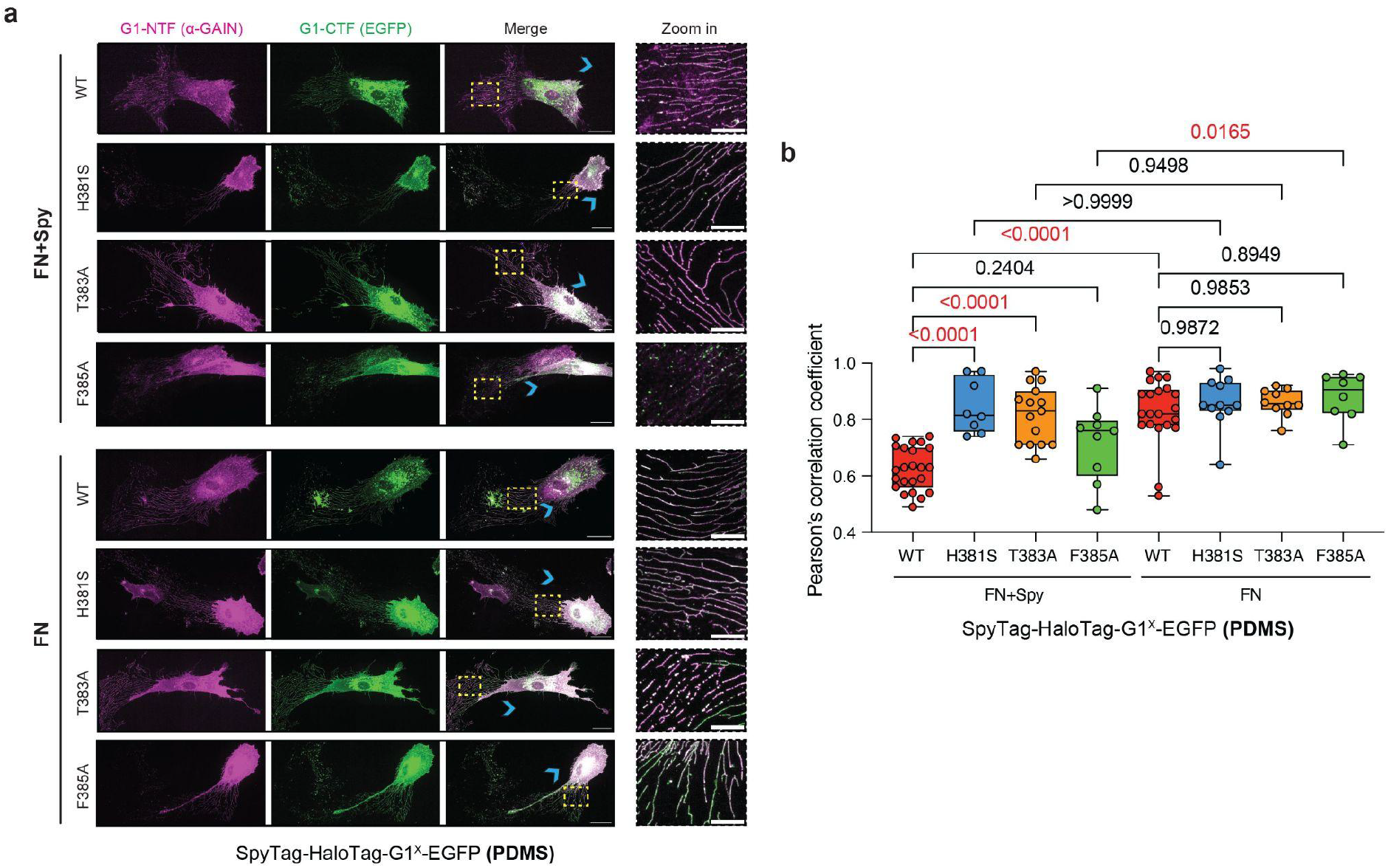
G1 dissociation of HFF cells with ‘migration force assays’. **(a)** Immunostaining of HFF cells expressing the indicated SpyTag-HaloTag-G1^x^-EGFP variants on FN+Spy or FN-coated surfaces, visualizing G1-NTF (α-GAIN) and G1-CTF (EGFP). Blue chevrons mark the direction of cell migration. Scale bars: 20 µm; zoom-in: 10 µm. **(b)** Pearson’s correlation coefficient between G1-NTF and G1-CTF signals in HFF cells. Data (n ≥ 8) are shown as box-and-whisker plots (median: horizontal line; boxes: 25th-75th percentiles; whiskers: min-max values). Ordinary one-way ANOVA with Tukey’s multiple comparisons test was performed; p-values are indicated.

**Supplementary Figure 4.**
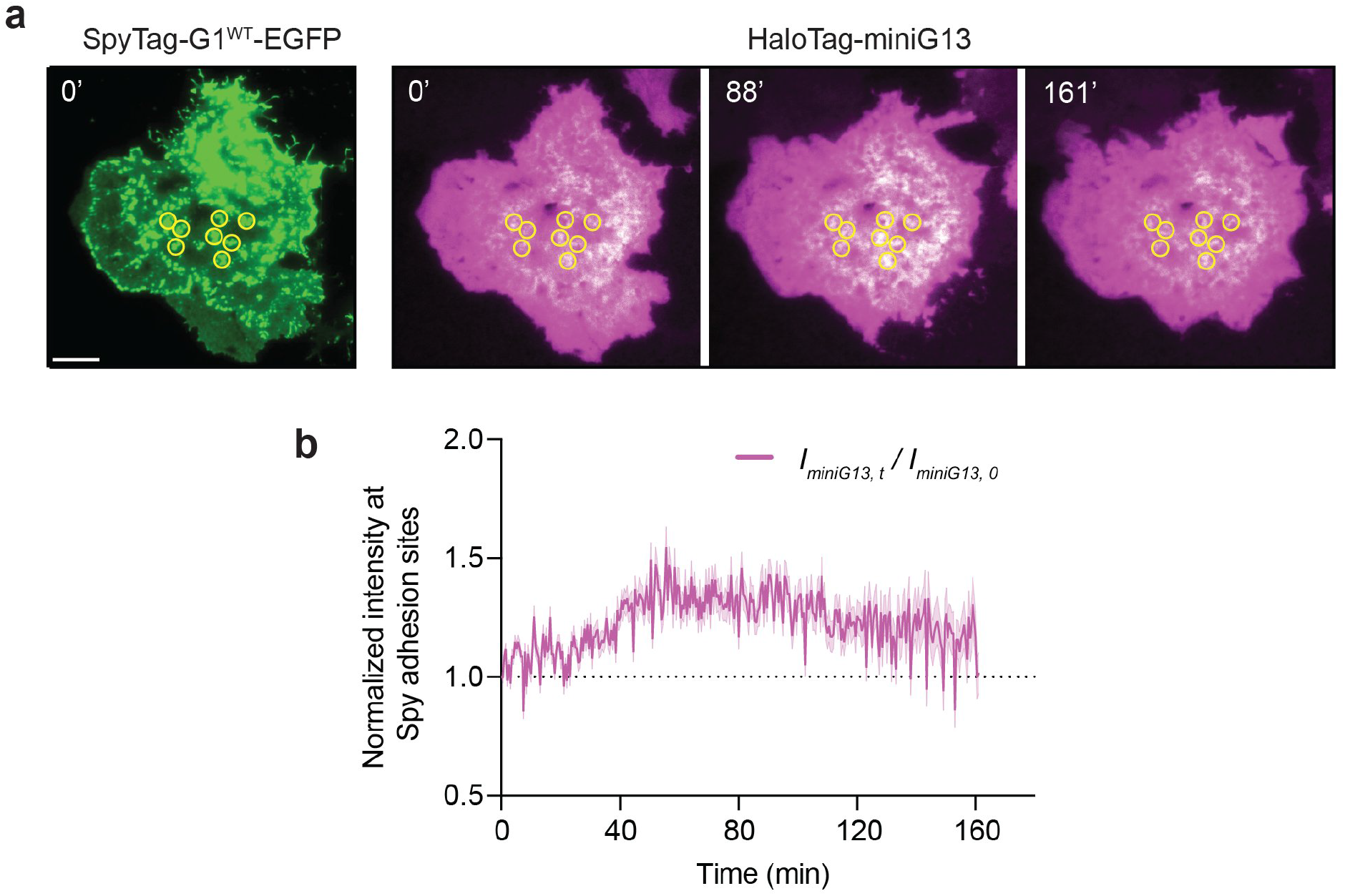
Saturation and decay of HaloTag-miniG13 signals during long term recording. **(a)** TIRFM images of cells expressing SpyTag-G1-EGFP (green) migrating on FN+Spy surfaces at indicated time points. HaloTag-miniG13 (magenta) is recruited to surface-expressed receptors in WT cells. HaloTag-miniG13 fluorescence increased and subsequently declined during prolonged imaging. Representative images are shown at 0, 88 and 161 min; the time trace is shown in **(b)**. Scale bar: 10 µm. **(b)** Dynamic changes in I_miniG13,t_/I_miniG13,0_ (magenta) at Spy adhesion sites over the 161-minute recording period.

**Supplementary Figure 5.**
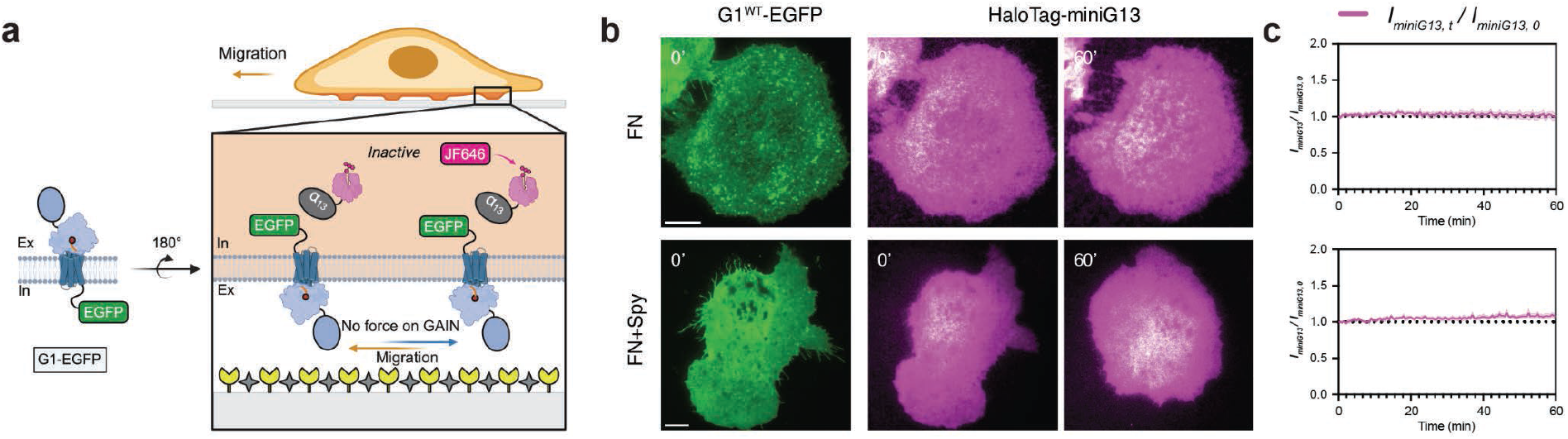
Requirement of adhesion site-formation or force generation through cell migration for G1 signaling and miniG13 recruitment. **(a)** G1-EGFP receptor design for the ‘migration force assay’, combining G protein recruitment imaging. No GAIN domain dissociation occurs at FN or FN+Spy surfaces, leading to no Halo-miniG13 recruitment. **(b-c)** TIRFM images of G1^WT^-EGFP expressing cells (green) migrating on FN or FN+Spy surfaces at indicated time points, with no HaloTag-miniG13 (magenta) recruitment over 60 minutes. Scale bar: 10 µm. Dynamic changes in I_miniG13,t_/I_miniG13,0_ (magenta) are shown; n ≥ 5 cells per group.

**Supplementary Figure 6.**
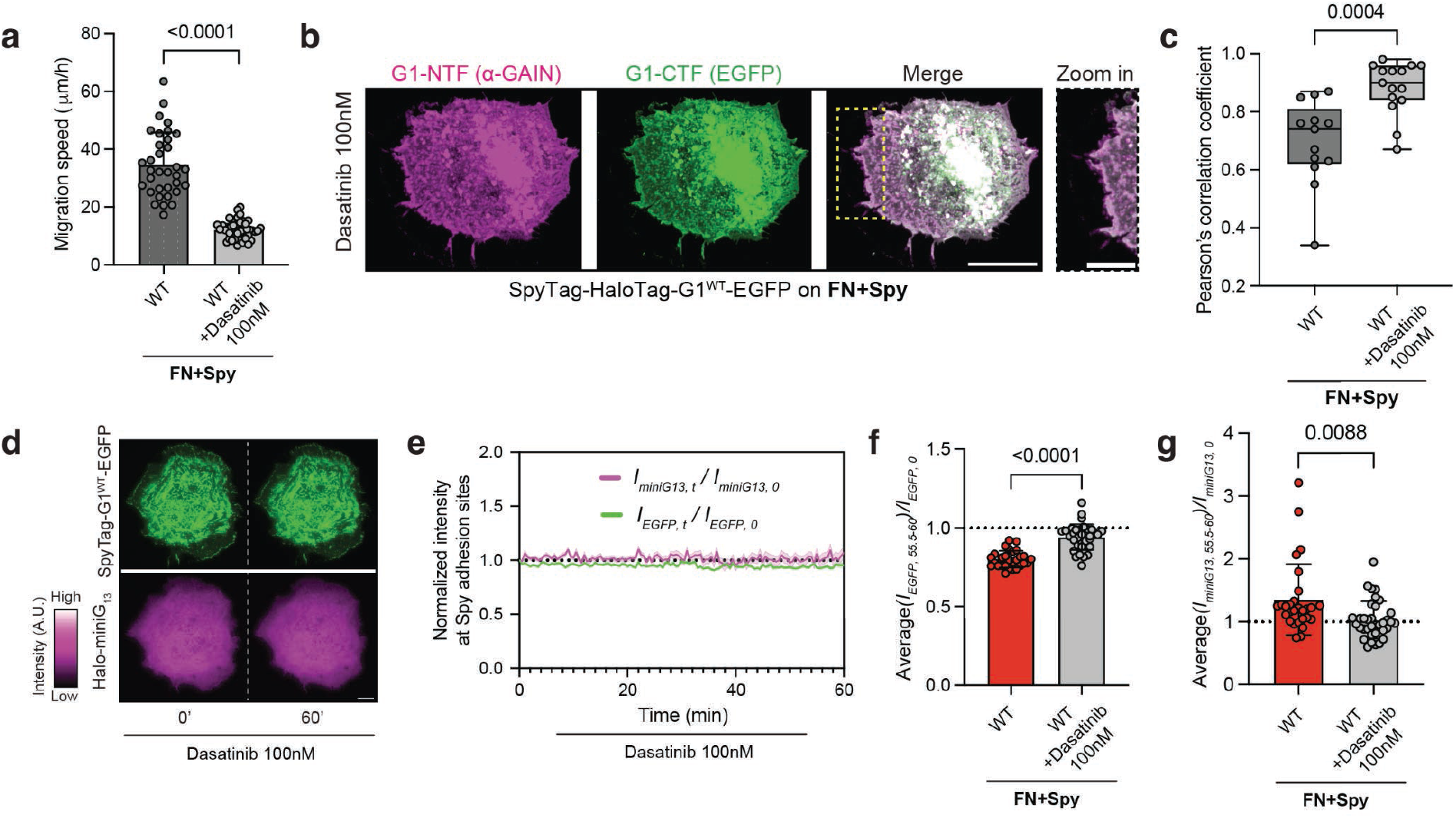
Cell migration is required for G1 dissociation, signaling and miniG13 recruitment. **(a)** Migration speed of COS-7 cells expressing SpyTag-HaloTag-G1-EGFP on FN+Spy surfaces with Dasatinib (100nM). Data (n ≥ 30 cells from 2 independent experiments) are shown as scatter plots with mean ± SD. Statistical analysis: Welch’s t-test (P values shown). **(b)** Immunostaining of COS-7 cells expressing SpyTag-HaloTag-G1-EGFP on FN+Spy surfaces with Dasatinib (100nM), visualizing G1-NTF (a-GAIN) and G1-CTF (EGFP). Scale bars: 20 µm; zoom-in: 10 µm. **(c)** Pearson’s correlation coefficient between G1-NTF and G1-CTF signals in COS-7 cells. Data (n ≥ 15 biological replicates from ≥ 2 independent experiments per group) are shown as box-and-whisker plots (median: horizontal line; boxes: 25th-75th percentiles; whiskers: min-max values). Welch’s t-test was performed; p-values are indicated. **(d)** TIRFM images of COS-7 cells expressing SpyTag-G1-EGFP (green) migrating on FN+Spy surfaces with Dasatinib (100 nM). No HaloTag-miniG13 (magenta) recruitment over 60 minutes. Scale bar: 10 µm. **(e)** Dynamic changes in normalized G1 intensity (I_EGFP,t_/I_EGFP,0_, green) and miniG13 intensity (I_miniG13,t_/I_miniG13,0_, magenta) at Spy adhesion sites over 60 minutes with Dasatinib (100 nM). **(f)** Average I_EGFP,t_/I_EGFP,0_ and **(g)** I_miniG13,t_/I_miniG13,0_ values from the final 5 minutes of imaging in **(e)**. Each point represents an adhesion site; at least 30 sites were sampled from at least six cells per group. Data are shown as scatter plots with mean ± SD. Statistical analysis: Welch’s t-test (P values shown). The same WT control dataset is shown in Suppl. Fig. 6f-g and 7d-e because the treatments were performed in parallel. Repeated display does not represent an additional independent experiment.

**Supplementary Figure 7.**
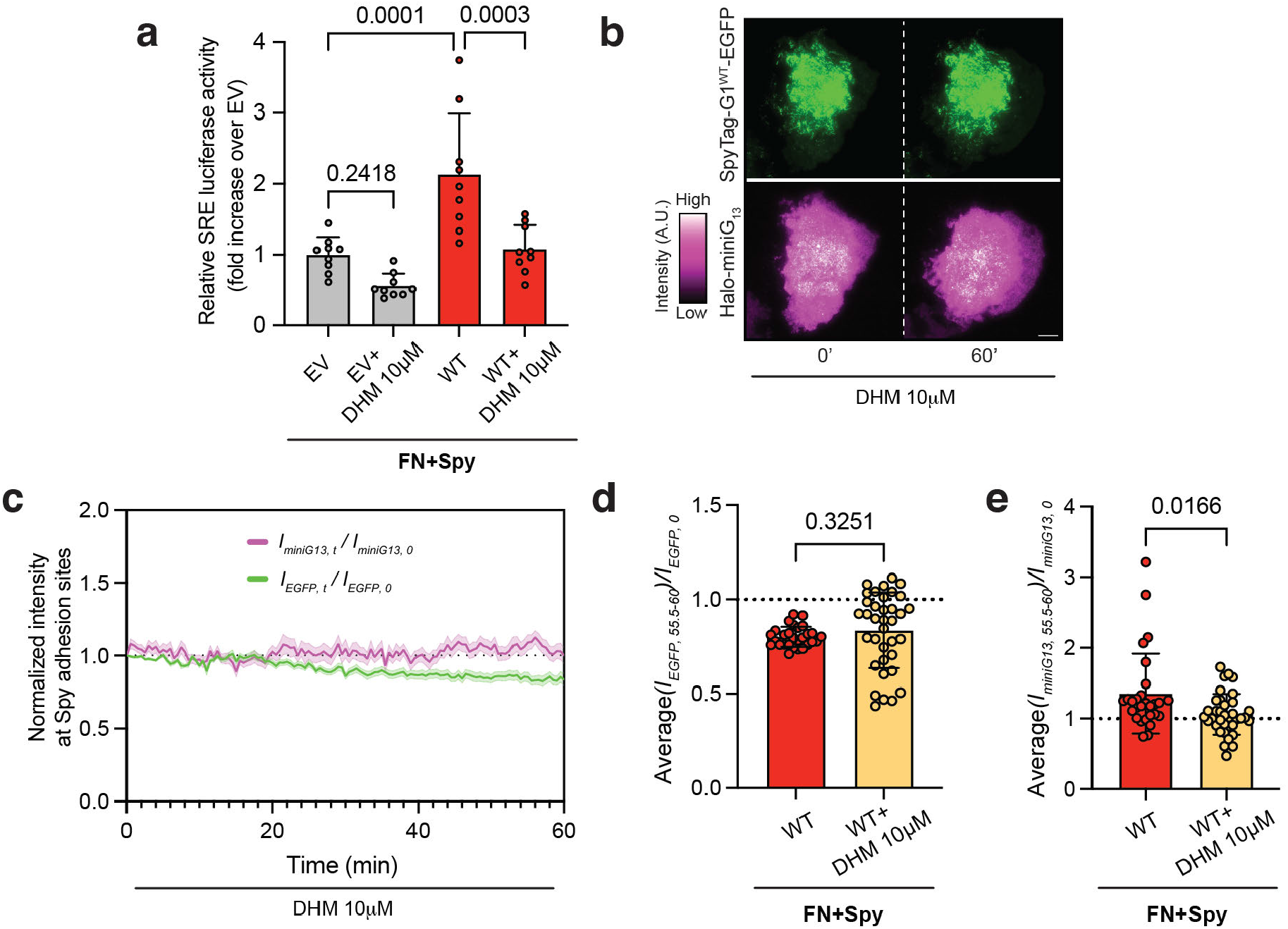
Migration-dependent force activation of G1 is mitigated by the partial G1 antagonist DHM. **(a)** Relative luciferase activity for G1 constructs with 10 µM DHM treatment on FN+Spy surfaces during migration. Data (n ≥ 3 replicates from 3 independent experiments) are shown as scatter plots with mean ± SD. Statistical analysis: one-way ANOVA with Tukey’s test (P values shown). **(b)** TIRFM images of COS-7 cells expressing SpyTag-G1-EGFP (green) migrating on FN+Spy surfaces with DHM (10 µM). No HaloTag-miniG13 (magenta) recruitment over 60 minutes. Scale bar: 10 µm. **(c)** Dynamic changes in normalized G1 intensity (I_EGFP,t_/I_EGFP,0_, green) and miniG13 intensity (I_miniG13,t_/I_miniG13,0_, magenta) at Spy adhesion sites over 60 minutes with 10 µM DHM treatment. **(d)** Average I_EGFP,t_/I_EGFP,0_ and **(e)** I_miniG13,t_/I_miniG13,0_ values from the final 5 minutes of imaging in **(c)**. 10 µM DHM treatment does not affect the G1 dissociation but reduces miniG13 recruitment. Each point represents an adhesion site; at least 30 sites were sampled from at least six cells per group. Data are shown as scatter plots with mean ± SD. Statistical analysis: Welch’s t-test (P values shown). The same WT control dataset is shown in Suppl. Fig. 6f-g and 7d-e because the treatments were performed in parallel. Repeated display does not represent an additional independent experiment.

**Supplementary Figure 8.**
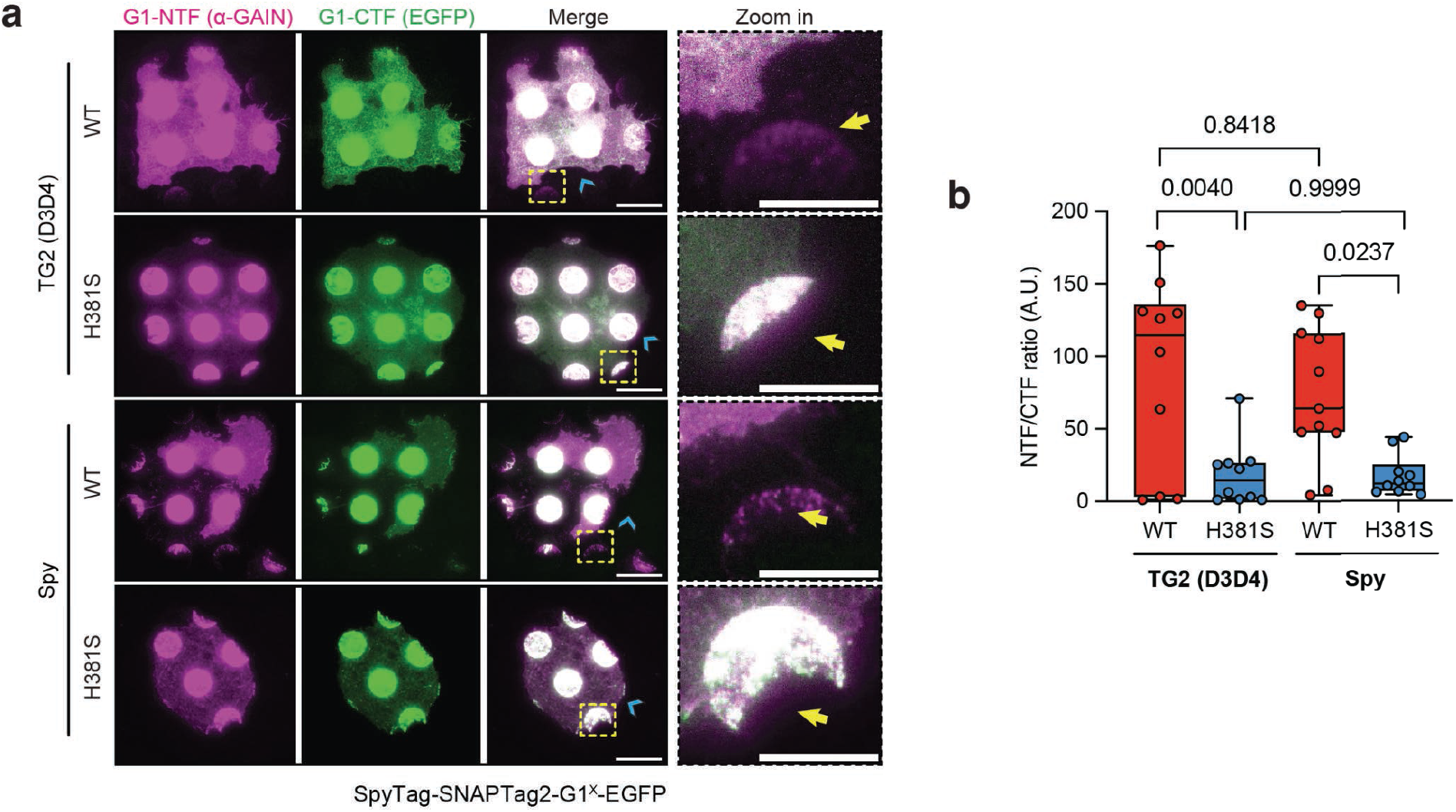
G1 dissociation occurs in WT but not H381S mutant on both Spy and TG2(D3D4)-coated micropatterns. **(a)** Immunostaining of COS-7 cells expressing the indicated SpyTag-SNAPTag2-G1^x^-EGFP variants on Spy and TG2 (D3D4)-coated micropatterns, visualizing G1-NTF (a-GAIN) and G1-CTF (EGFP). Scale bars: 20 µm; zoom-in: 10 µm. **(b)** NTF/CTF ratio for indicating G1 dissociation in **(a)**. Data (n ≥ 10 biological replicates from ≥ 2 independent experiments per group) are shown as box-and-whisker plots (median: horizontal line; boxes: 25th-75th percentiles; whiskers: min-max values). Ordinary one-way ANOVA with Tukey’s multiple comparisons test was performed; p-values are indicated.

**Supplementary Figure 9.**
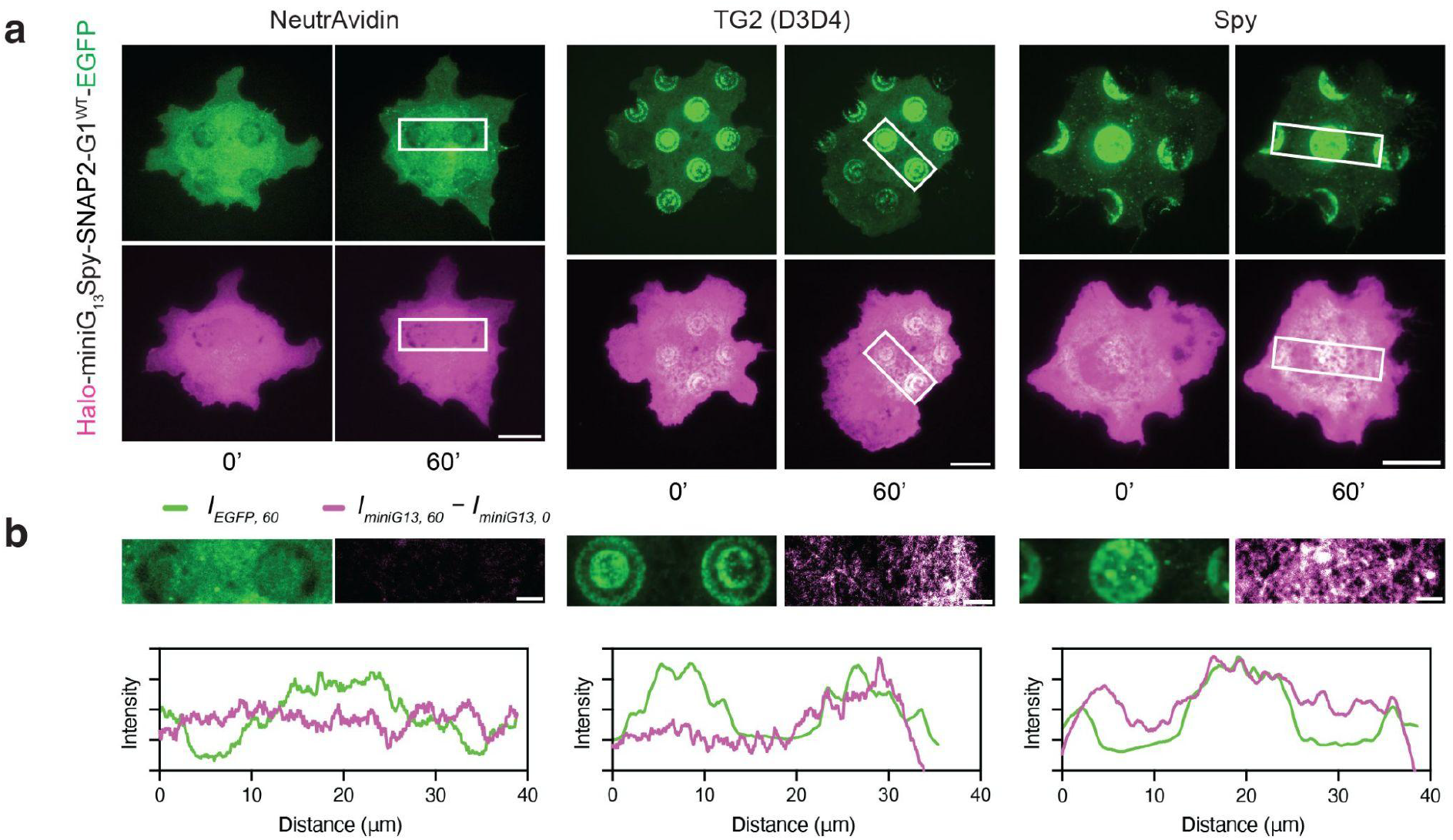
Mechano-dependent G1 activation at TG2 (D3D4)- and Spy-coated micropatterns during cell migration. **(a)** TIRFM images of COS-7 cells expressing SpyTag-SNAPTag2-G1^WT^-EGFP(green) migrating on uncoated (left), TG2 (D3D4)-coated (middle), and SpyCather-coated (right) micropatterns. G1 forms adhesions on both TG2 (D3D4) and SpyCatcher micropatterns. HaloTag-miniG13 (magenta) is recruited to WT G1 on these micropatterns. A zoomed-in white box (∼40 µm, shown in **(b)**) highlights the 60-minute time point. Scale bar: 20 µm. **(b)** Zoom-in region (top) and normalized intensity profiles (bottom) for G1 (green) and miniG13 (magenta) signal at uncoated (left), TG2 (D3D4)-coated (middle), and SpyCather-coated (right) micropatterns. MiniG13 intensity reflects net increases during the 60-minute interval. Scale bar: 5 µm.

**Supplementary Figure 10.**
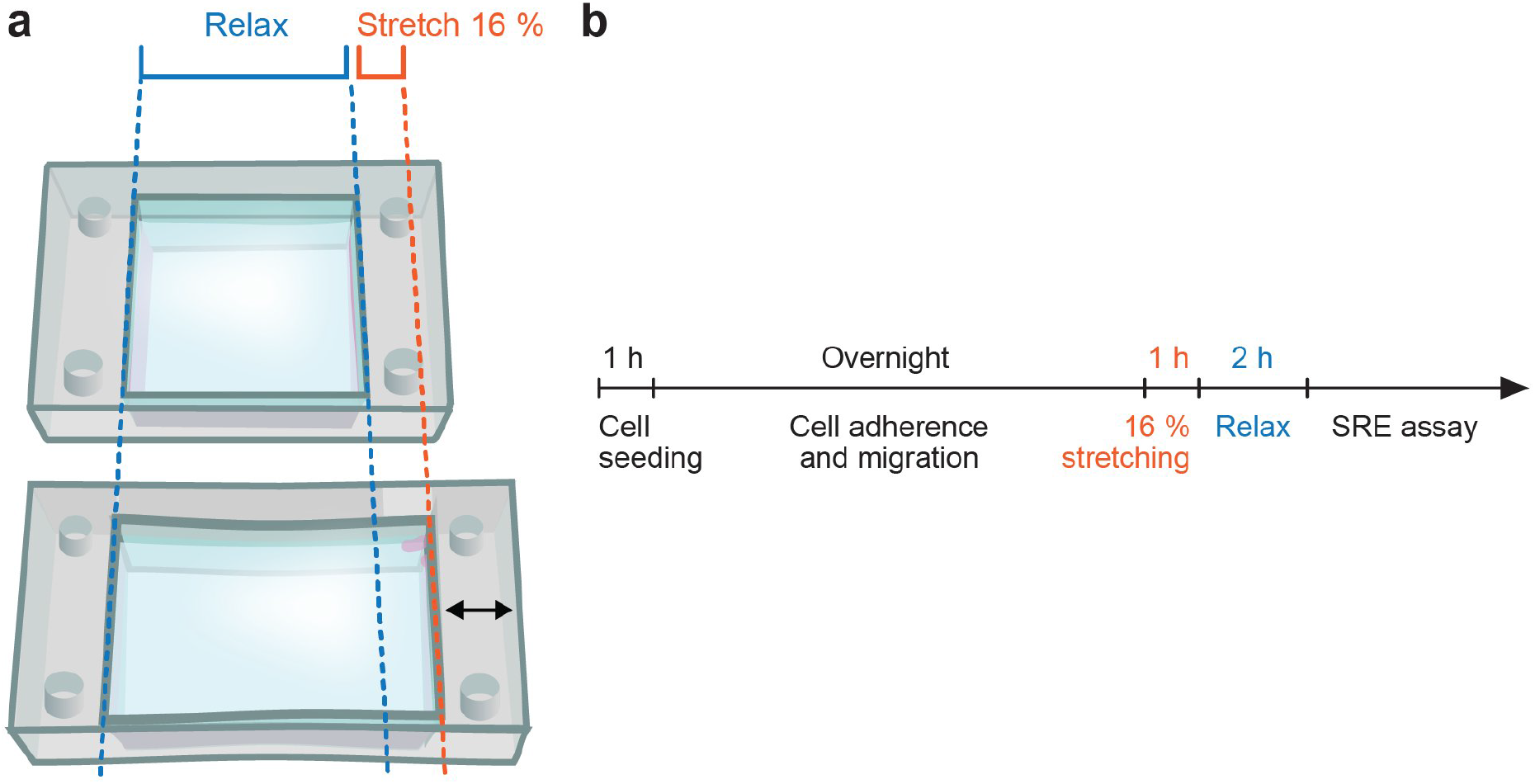
Schematic of the stretching chamber and the protocol for SRE measurement for the ‘deformation force assay’. **(a)** Schematic of the stretching chamber for the ‘deformation force assay’. COS-7 cells with the indicated constructs were seeded on FN- or FN+Spy-coated chambers. Substrates were either left unstretched (top) or stretched by 16 % (bottom). **(b)** Protocol for SRE measurement following the ‘deformation force assay’.

**Supp. Table S1.**
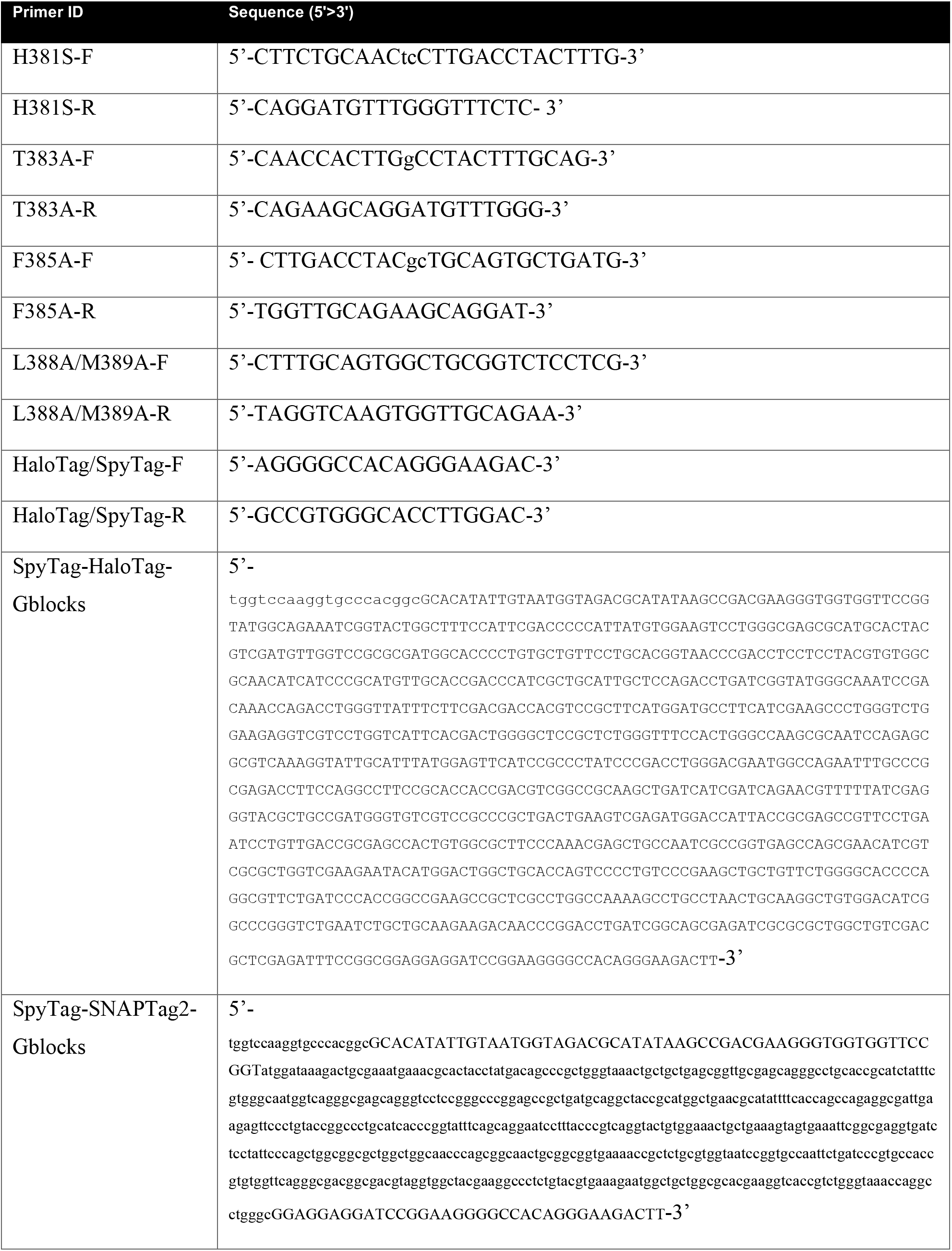
Primers used in this study.

**Supp. Video S1.**

Migration of COS-7 with respective SpyTag-HaloTag-G1-EGFP constructs on FN+Spy surface, recorded at 1 frame/min. Scale bar, 10 µm.

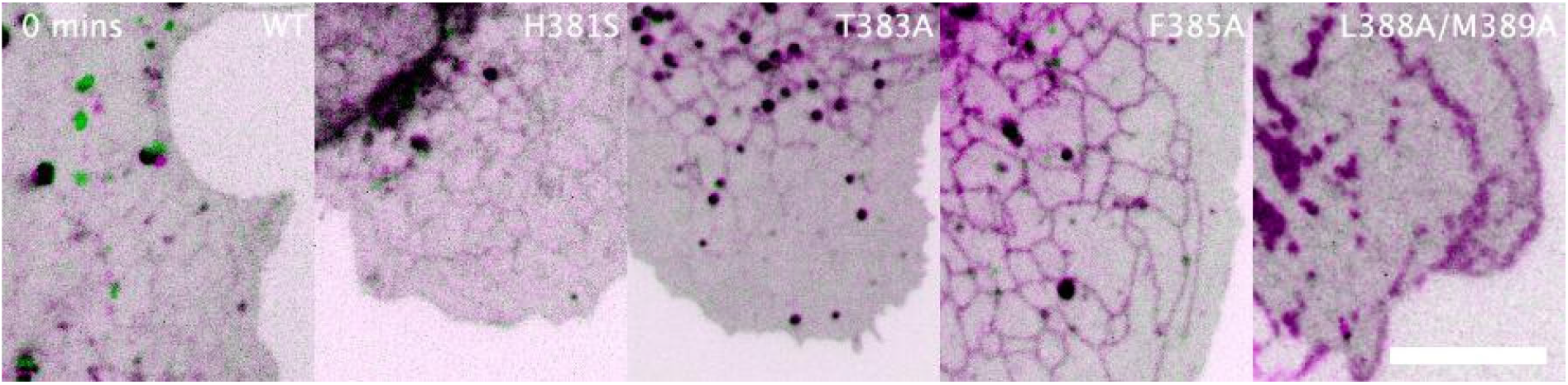

**Supp. Video S2.**

Migration of COS-7 with SpyTag-G1-EGFP and HaloTag-miniG13 constructs on FN+Spy surface, recorded at 2 frame/min. 1 hour duration. Scale bar, 10 µm.

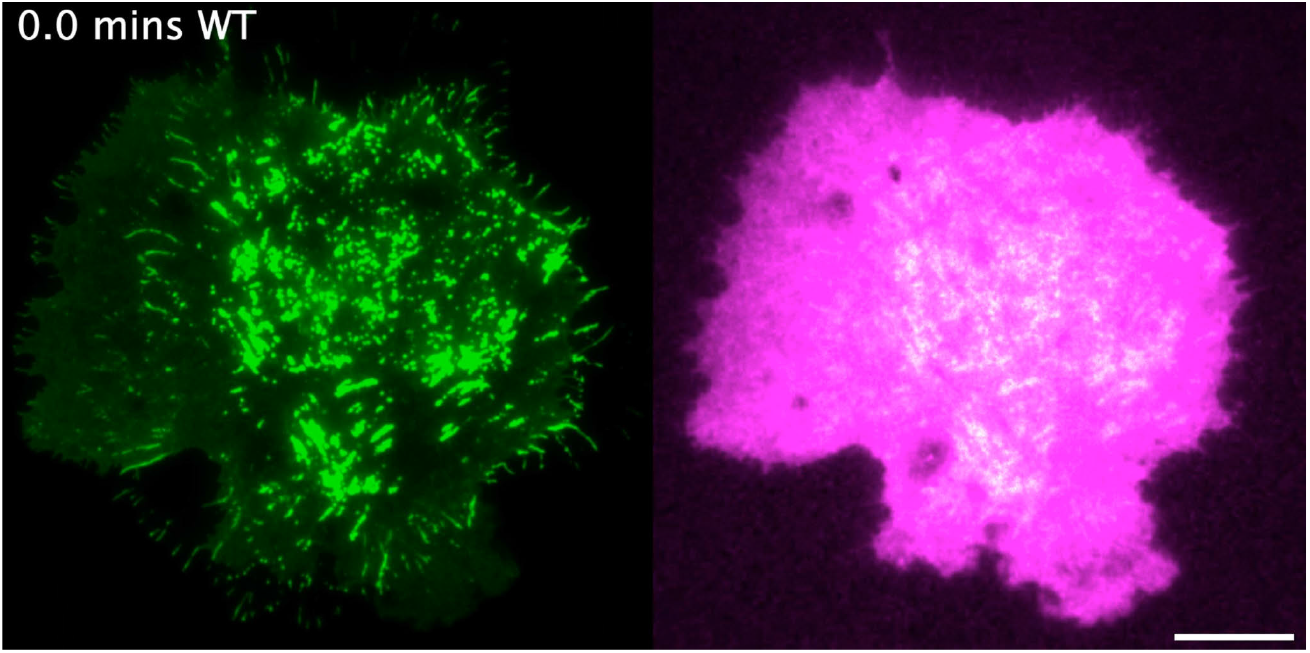

**Supp. Video S3.**

Migration of COS-7 with SpyTag-G1-EGFP and HaloTag-miniG13 constructs on FN+Spy surface, recorded at 2 frame/min. 161 minutes duration. Scale bar, 10 µm.

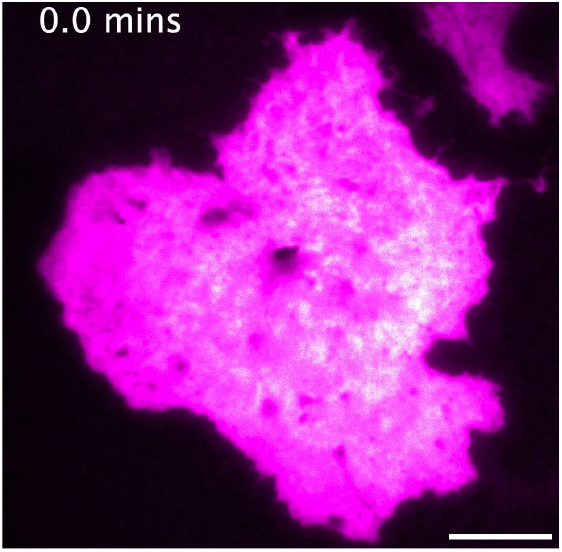

**Supp. Video S4.**

Migration of COS-7 with SpyTag-G1^381S^-EGFP and HaloTag-miniG13 constructs on FN+Spy surface, recorded at 2 frame/min. 1 hour duration. Scale bar, 10 µm.

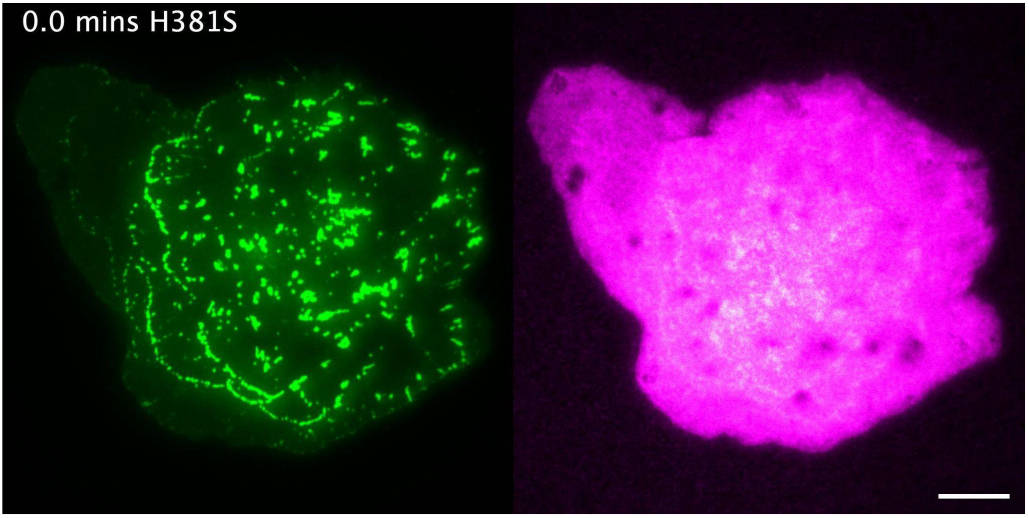

**Supp. Video S5.**

Migration of COS-7 with SpyTag-G1^T383A^-EGFP and HaloTag-miniG13 constructs on FN+Spy surface, recorded at 2 frame/min. 1 hour duration. Scale bar, 10 µm.

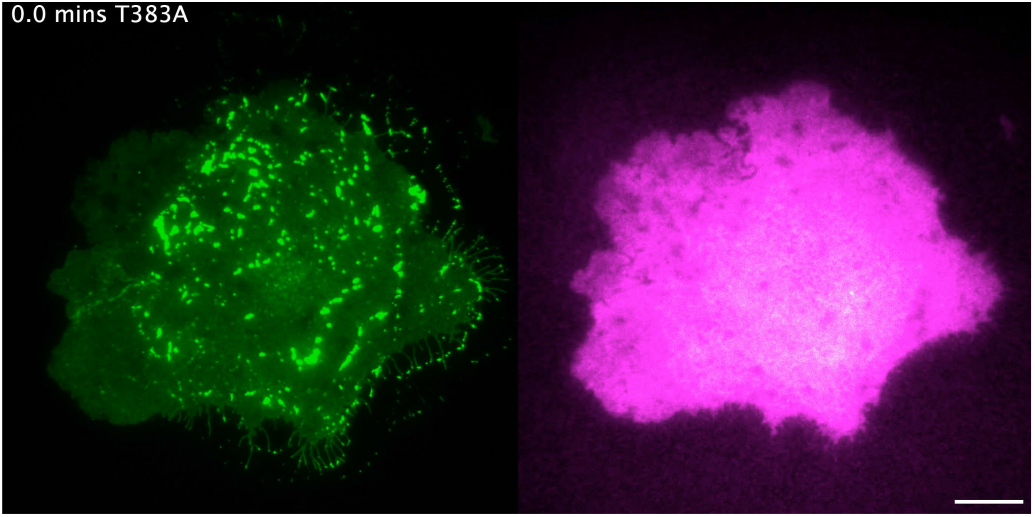

**Supp. Video S6.**

Migration of COS-7 with SpyTag-G1^F385A^-EGFP and HaloTag-miniG13 constructs on FN+Spy surface, recorded at 2 frame/min. 1 hour duration. Scale bar, 10 µm.

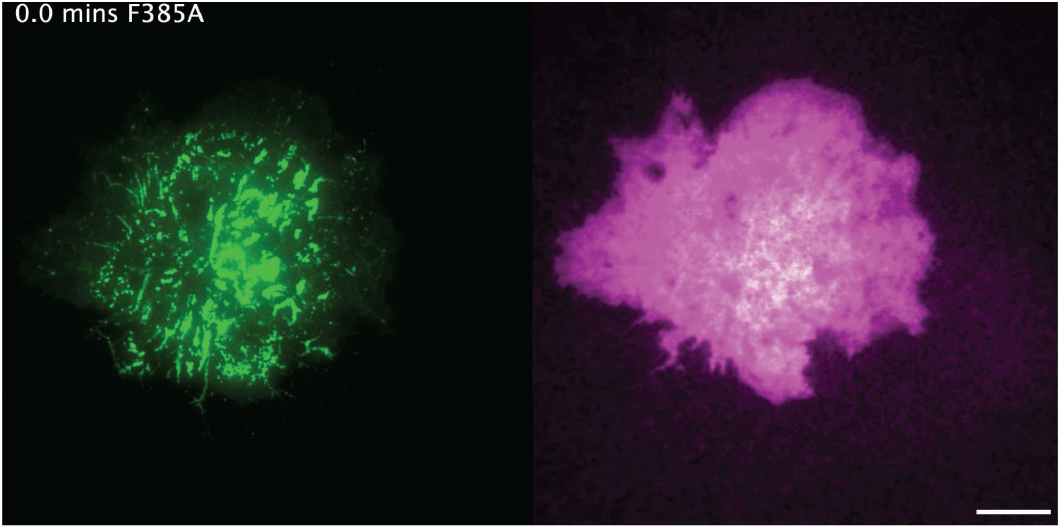

**Supp. Video S7.**

Migration of COS-7 with SpyTag-G1^L388A/M389A^-EGFP and HaloTag-miniG13 constructs on FN+Spy surface, recorded at 2 frame/min. 1 hour duration. Scale bar, 10 µm.

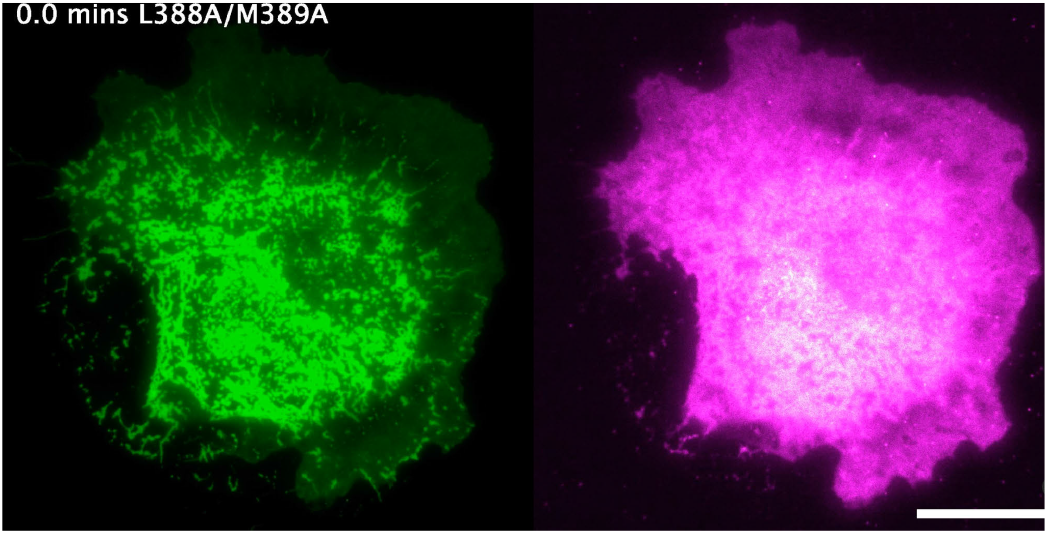

**Supp. Video S8.**

Zoom-in region for G1 (green) and miniG13 (magenta) signal at Spy adhesion sites. MiniG13 intensity reflects net increases during the 60-minute interval, recorded at 2 frame/min. 1 hour duration. Scale bar: 3 µm.

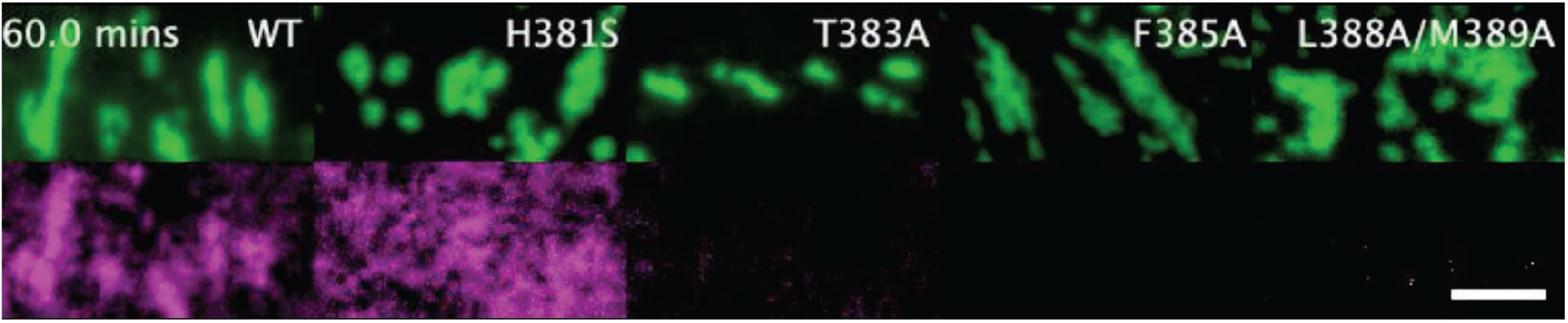

**Supp. Video S9.**

Migration of COS-7 with SpyTag-SNAPTag2-G1-EGFP and HaloTag-miniG13 constructs on TG2(D3D4)-coated micropatterns with FN in the surrounding area, recorded at 2 frame/min. 1 hour duration. Scale bar, 20 µm.

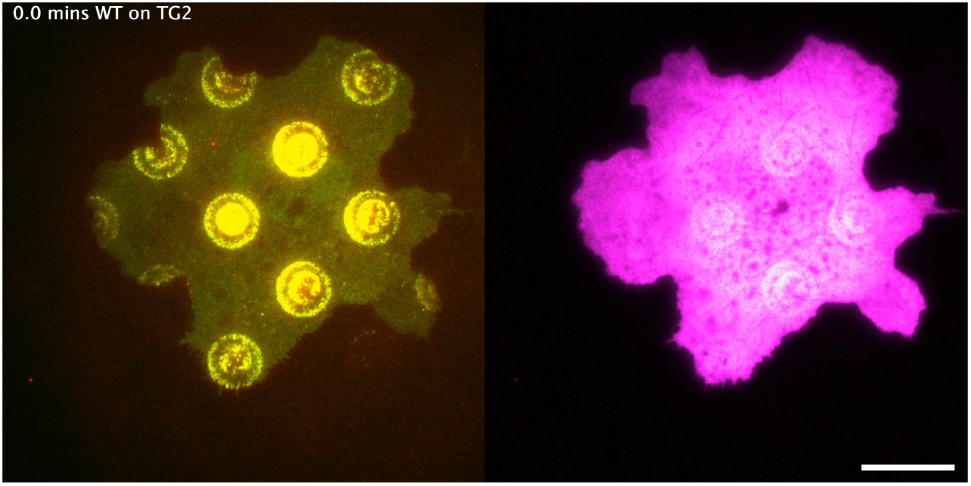

**Supp. Video S10.**

Zoom-in region (12 µm in diameter) showing G1 dissociation (top) and miniG13 recruitment (bottom) at TG2(D3D4)-coated micropatterns, recorded at 2 frame/min. 1 hour duration. Scale bar, 5 µm.

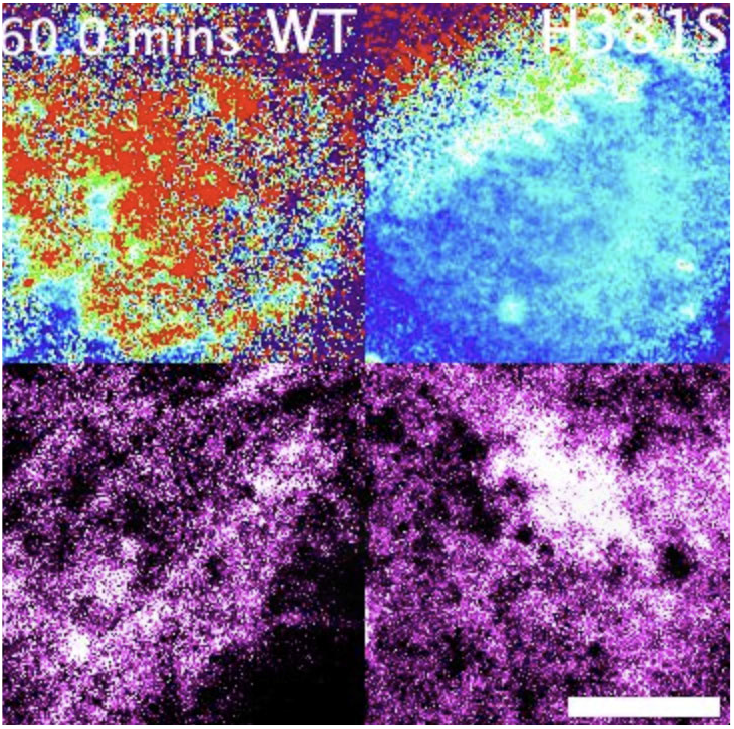

**Supp. Video S11.**

Migration of COS-7 with SpyTag-SNAPTag2-G1^H381S^-EGFP and HaloTag-miniG13 constructs on TG2(D3D4)-coated micropatterns with FN in the surrounding area, recorded at 2 frame/min. 1 hour duration. Scale bar, 20 µm.

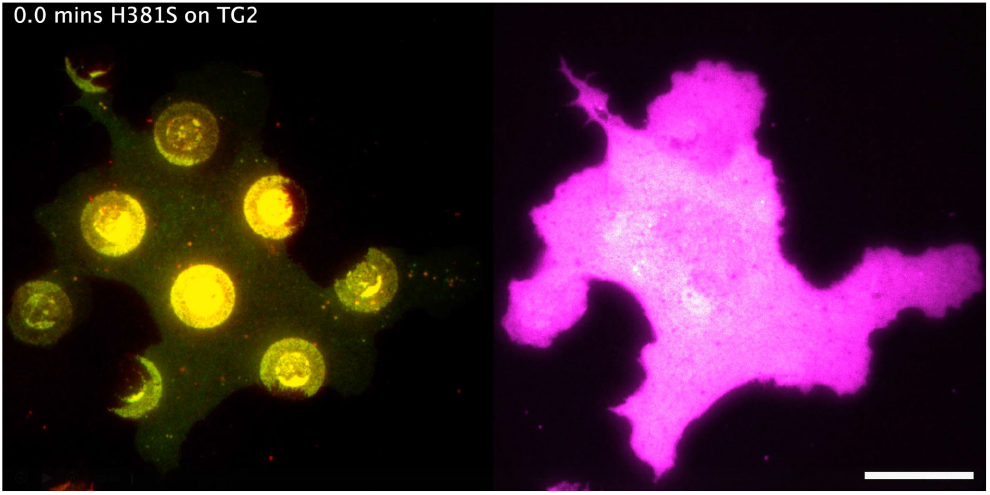

**Supp. Video S12.**

Migration of COS-7 with SpyTag-SNAPTag2-G1^T383A^-EGFP and HaloTag-miniG13 constructs on TG2(D3D4)-coated micropatterns with FN in the surrounding area, recorded at 2 frame/min. 1 hour duration. Scale bar, 20 µm.

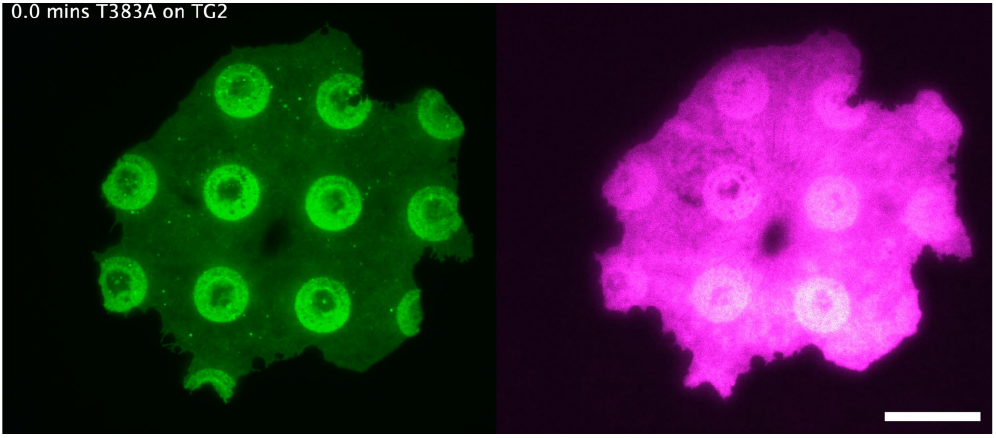

**Supp. Video S13.**

Migration of COS-7 with SpyTag-SNAPTag2-G1^F385A^-EGFP and HaloTag-miniG13 constructs on TG2(D3D4)-coated micropatterns with FN in the surrounding area, recorded at 2 frame/min. 1 hour duration. Scale bar, 20 µm.

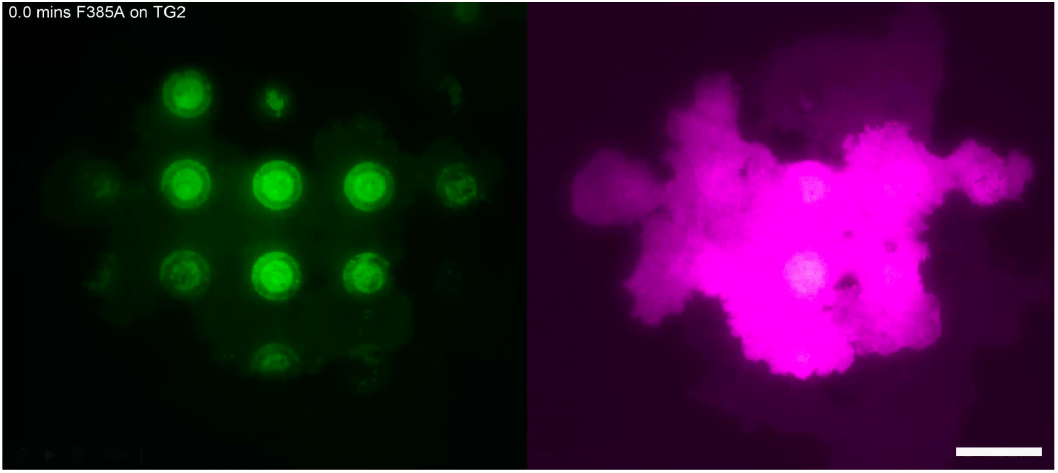

**Supp. Video S14.**

Migration of COS-7 with SpyTag-SNAPTag2-G1^L388A/M389A^-EGFP and HaloTag-miniG13 constructs on TG2(D3D4)-coated micropatterns with FN in the surrounding area, recorded at 2 frame/min. 1 hour duration. Scale bar, 20 µm.

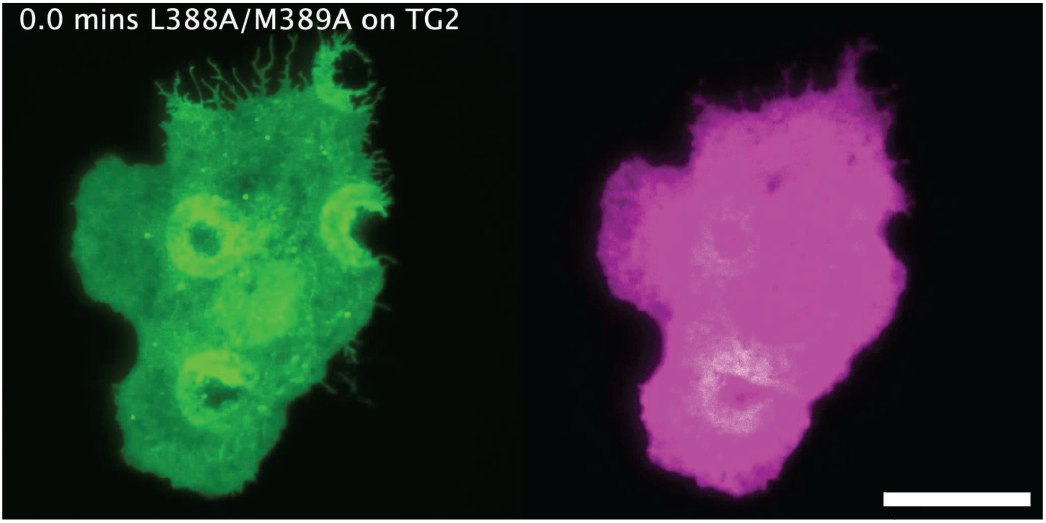

